# Genome-wide approaches delineate the additive, epistatic, and pleiotropic nature of variants controlling fatty acid composition in peanut (*Arachis hypogaea L*.)

**DOI:** 10.1101/2021.06.03.446924

**Authors:** Paul I. Otyama, Kelly Chamberlin, Peggy Ozias-Akins, Michelle A. Graham, Ethalinda K. S. Cannon, Steven B. Cannon, Gregory E. MacDonald, Noelle L. Anglin

**Affiliations:** Interdepartmental Genetics and Genomics, Iowa State University, Ames, IA; Agronomy Department, Iowa State University, Ames, IA; ORISE Postdoctoral Fellow, Corn Insects and Crop Genetics Research Unit, USDA-ARS, Ames, IA; USDA - Agricultural Research Service, Stillwater, OK; Institute of Plant Breeding, Genetics, and Genomics and Department of Horticulture, University of Georgia, Tifton, GA; USDA - Agricultural Research Service, Corn Insects and Crop Genetics Research Unit, Ames, IA; University of Florida, Gainesville, FL; USDA-ARS Small Grains and Potato Research Laboratory, Aberdeen, ID

**Author notes:** co-senior authors (Gregory E. MacDonald, Noelle L. Anglin). **Author for correspondence**: Steven B. Cannon: 1017 Crop Genome Informatics Laboratory, 819 Wallace Rd, Ames, Iowa 50011.

## Abstract

The fatty acid composition of seed oil is a major determinant of the flavor, shelf-life, and nutritional quality of peanuts. Major QTLs controlling high oil content, high oleic content, and low linoleic content have been characterized in several seed oil crop species. Here we employ genome-wide association approaches on a recently genotyped collection of 787 plant introduction accessions in the USDA peanut core collection, plus selected improved cultivars, to discover markers associated with the natural variation in fatty acid composition, and to explain the genetic control of fatty acid composition in seed oils.

Overall, 251 single nucleotide polymorphisms (SNPs) had significant trait associations with the measured fatty acid components. Twelve SNPs were associated with two or three different traits. Of these loci with apparent pleiotropic effects, 10 were associated with both oleic (C18:1) and linoleic acid (C18:2) content at different positions in the genome. In all 10 cases, the favorable allele had an opposite effect - increasing and lowering the concentration, respectively, of oleic and linoleic acid. The other traits with pleiotropic variant control were palmitic (C16:0), behenic (C22:0), lignoceric (C24:0), gadoleic (C20:1), total saturated, and total unsaturated fatty acid content. One hundred (100) of the significantly associated SNPs were located within 1000 kbp of 55 genes with fatty acid biosynthesis functional annotations. These genes encoded, among others: ACCase carboxyl transferase subunits, and several fatty acid synthase II enzymes.

With the exception of gadoleic (C20:1) and lignoceric (C24:0) acid content, which occur at relatively low abundance in cultivated peanut, all traits had significant SNP interactions exceeding a stringent Bonferroni threshold (**α** = 1%). We detected 7,682 pairwise SNP interactions affecting the relative abundance of fatty acid components in the seed oil. Of these, 627 SNP pairs had at least one SNP within 1000 kbp of a gene with fatty acid biosynthesis functional annotation. We evaluated 168 candidate genes underlying these SNP interactions. Functional enrichment and protein-to-protein interactions supported significant interactions (p- value < 1.0E-16) among the genes evaluated. These results show the complex nature of the biology and genes underlying the variation in seed oil fatty acid composition and contribute to an improved genotype-to-phenotype map for fatty acid variation in peanut seed oil.

**Key phrases:** SNP Genotyping, Genome-wide Association Study (GWAS), GWAS of interacting SNPs (GWASi), Pleiotropy, Seed fatty acid composition, Oleic-Linoleic acid ratio.

## Introduction

Cultivated peanut or groundnut (*Arachis hypogaea* L.) is a food and oil crop of global importance. Dry peanut seeds are ∼46 – 58% oil and as high as 22 – 32% protein (Dean et al. 2009; Dezern 2018). Peanut oil is enriched with antioxidants and heart-healthy unsaturated fats. Thus, it ranks fifth in global production and consumption after palm, soybean, rapeseed, and sunflower. Annual global production is approximately 47 M tons of unshelled nuts, yielding about 5.8 M metric tons of seed oil. China and India account for ∼70% of the global production, followed by Africa and the Americas at 22% and 4.1% (http://www.fao.org/faostat/en/#data/QD, accessed 01/04/2021 at 4:50 pm CST). The fatty acid profile of peanut oil is an important seed quality trait. The oil is composed of varying amounts of both saturated and unsaturated fats. Long chain fatty acids are synthesized de novo from photosynthetic precursors in composite reaction steps catalyzed by acetyl- Coenzyme A carboxylase (ACCase) and fatty acid synthase (FAS) enzyme systems. Acyl carrier proteins (ACP) are important cofactors in this process. In plants, FAS enzymes function as discrete units catalyzing single reactions (FAS II enzyme system), while in animals and fungi, FAS enzymes exist as multifunctional polypeptides with domains for individual activities (type I FAS) (Chen et al. 2011; Gunstone and Harwood 2007; Harwood 2005; Phan et al. 2015).

In plants, de novo synthesis is initiated by the carboxylation of acetyl-CoA to form malonyl-CoA. This reaction is catalyzed by ACCase in the plastids. What follows is a four-step elongation cycle, during which the fatty acid chain is extended by two carbons in each round. Ketoreductases (KR), dehydratases (DH), and enoyl reductases (ER) are the active enzymes during this stage, catalyzing the reduction of the beta keto group to a fully saturated carbon chain. Synthesis is completed after six recurring reactions with the formation of a 16-carbon long chain saturated fatty acid, palmitate [16:0] (Chen et al. 2011; Harwood 2005).

The synthesized palmitate undergoes several modifications starting with elongation to form stearic acid (18:0). These two are the most abundant saturated fatty acids in peanuts and many plant oils. Elongase enzymes catalyze reactions producing very long chain fatty acids (>18C) from palmitate and stearate. Desaturase enzymes catalyze reactions producing unsaturated fatty acids from palmitate, stearate, and/or very long chain fatty acids. Desaturation involves the introduction of double bonds in one or several positions along the fatty acyl chain (Chen et al. 2011; Phan et al. 2015).

Many fatty acid desaturase genes (FAD) have been identified and characterized in various oil crops. FADs are named and numbered according to the position of the double bond introduced in the chain, as well as on the nature of their substrate. Stearoyl-ACP Δ9-desaturase, a soluble enzyme in the plastid stroma, converts stearate (18:0) to oleate (18:1). Δ12-desaturase, encoded by FAD2, introduces methylene-interrupted double bond arrangement into oleate to produce linoleate (18:2) which can then be converted into α-linolenate by a Δ15-desaturase enzyme encoded by FAD3. Further modification of straight chain fatty acids occurs via the addition of Epoxy, Oxy, Hydroxy, and Acetylenic groups to form cyclic fatty acids which have many Industrial applications (Chen et al. 2011; Phan et al. 2015).

Unlike FAD2 and Stearoyl-ACP Δ9-desaturase genes, which have been well studied and characterized across several plant species, many of the other genes responsible for the abundance and variation of fatty acids in seed oils remain understudied, along with their genetic controls. We leveraged a large, recently genotyped collection of diversely sourced, unimproved peanut accessions, as well as selected improved cultivars (Otyama et al. 2020), to study the variation in seed oil fatty acid composition. We employed genome-wide association approaches to identify regions in the genome – both novel and previously studied - that act additively, in concert, and pleiotropically, to explain the genetic control of fatty acid composition in peanut seed oils.

## Materials and Methods

### Plant Material and Field Layout

The experiment was planted in replicates over two years, in Citra, Florida, at the University of Florida Plant Science Research and Education Unit. Materials were planted in 2013, between May 12 – 15, and in 2015, between April 27 – 30. An Augmented Randomized Block design with three blocks was used. Fourteen (14) improved cultivars which included high and normal oleic peanut cultivars were planted in each of the three blocks as checks. Of these, eight were replicated three times and the remaining six were replicated once (due to seed shortages) in each block. 687 lines representing the US peanut core collection were planted as treatments split randomly among the three blocks. Additionally, 107 mini core lines (a subset of the core) were replicated once in each block. Seed biochemical composition was assayed as described in our previous work for total oil, total protein, and fatty acid components of seed oil (Dezern 2018; Otyama et al. 2019). The following fatty acids were measured: palmitic (C16:0), stearic (C18:0), oleic (C18:1), linoleic (C18:2), arachidic (C20:0), gadoleic (C20:1), behenic (C22:0), and lignoceric (C24:0).

### Statistical Analysis

To estimate best linear unbiased predictions (BLUPs), linear models were implemented in R using the function ***lmer*** from the package ***lme4*** (Bates et al. 2007) which exploits REML to estimate individual BLUPs and Variance components using the model:

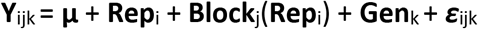

Where: **Y**_ijk_ is the trait of interest, **µ** is the mean effect, **Rep**_i_ is the effect of the **i**th replicate, **Block**_j(_**Rep**_i_ is the effect of the **j**th incomplete block within the **i**th replicate, **Gen**_k_ is the effect of the **k**th genotype, and ***ε***_ijk_ is the error associated with the **i**th replication, **j**th incomplete block and the **k**th genotype, which is assumed to be normally and independently distributed, with mean zero and homoscedastic variance **σ**^2^.

Broad-sense heritability of a given trait at an individual environment was calculated as:

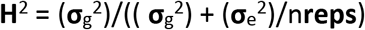

Where: **σ**_g_^2^ and **σ**_e_^2^ are the genotype and error variance components, and n**reps** is the number of replicates. The estimated broad-sense heritability is a measure of repeatability for each trait value – given that only a fraction of the genotypes was replicated due to the large size of the collection. All effects are considered random for calculating BLUPs and broad-sense heritability. Only one year of data was available for this study. The BLUP phenotypic distributions for each trait were plotted to check normality assumptions.

### Genotyping and Quality control

Accessions and cultivars were genotyped as described in Otyama et al. (2020) which yielded a total of 14,430 high-confidence SNPs, for the 1,120 *Arachis* samples. To test for the effect of population size and allelic diversity on high quality trait associations, first – 885 unique accessions and cultivars were selected based on a 98% identity threshold from all genotyped samples (Otyama et al. 2020); and for these, 13,410 markers were polymorphic. Of these unique subjects, three sub-populations were selected at 75%, 50%, and 25% selection intensities for minimum repetitiveness and maximum allelic diversity within the selected groups. We used the tool *Core Hunter II* (De Beukelaer et al. 2012) to sample and select, resulting in population subsets with 664, 442, and 221 subjects, respectively. Prior to GWAS, SNPs from each subpopulation were filtered for less than 30% missing data, minimum 5% minor allele frequency (MAF), and less than 20% heterozygosity. The remaining missing positions were then imputed using *Link Impute* in *TASSEL 5* (Bradbury et al. 2007; Money et al. 2015) to yield a final set of 11,038, 11,017, and 11,052 SNP markers for GWAS, with 664, 442, and 221 subjects, respectively.

### Single SNP Association Analysis (GWAS)

Single SNP associations were tested for each trait using the Fixed and random model Circulating Probability Unification model (FarmCPU) in the Genomic Association and Prediction Integrated Tool (Lipka et al. 2012; Liu et al. 2016). Stratification in the data was assessed using principal component analysis (PCA) and controlled by fitting different models with varying levels of control ranging from 0, where no principal components were added to the model, to 4, where 4 PCs were fitted to account for structure. The extent of model fitting was confirmed using a quantile–quantile (Q-Q) plot for the expected and obtained association p-values.

Resulting p-values were corrected using a modified Bonferroni adjustment based on linkage disequilibrium (LD) pruning to determine the most significant associations. We defined the number of independent statistical tests as the sum of the independent LD blocks plus singleton markers, which resulted in 3249 independent tests (Gao et al. 2010; Gao et al. 2008). We thus constructed two significance thresholds: 1) a 5% genome-wide significance threshold (0.05/3249, 1.54E-05), and 2), a more stringent genome-wide threshold (0.001/3249, 3.08E-07). LD was calculated in *TASSEL 5* (Bradbury et al. 2007) and decay was plotted in R as described in Otyama et al. (2019) using the Hill and Weir method, later modified by Remington (Hill and Weir 1988; Remington et al. 2001).

A window of 1000 kbp around each significant SNP – corresponding to an r^2^ of ∼0.35, was considered to obtain an overlapping set of candidate genes for each trait. The window size was chosen based on the genomic linkage disequilibrium of this peanut collection, which is quite high compared to other self-pollinating species. To further explore the results, all predicted gene models within the window were extracted for enrichment analysis. The quantile–quantile (Q–Q) plot and Manhattan plots of GWAS results were produced using the package qqman and CMplot in R (Turner 2014; Yin et al. 2021).

### SNP-SNP Interaction Association Analysis (GWASi)

Due to epistasis, two or more genes can interact to contribute to a single phenotype in a non-additive manner. To determine the contribution of SNP interactions to the overall variation in seed oil fatty acid composition, genome-wide SNP interactions were tested for each fatty acid component.

We implemented two strategies to screen for GWASi: an exhaustive screen, where all possible pairwise interactions across all genome-wide SNPs were tested; and a selective approach, testing pairwise interactions across select SNPs based on their proximity (1000 kbp) to genes annotated for their role in fatty acid biosynthesis. We used Tifrunner gene annotations (Tifrunner genome 2 annotation 1), accessed through the PeanutBase data store on PeanutBase We also included SNPs with significant trait associations from GWAS (P values < 1.54E-05) in our selective testing approach. In both GWASi strategies, markers were subjected to LD pruning to limit top ranked interactions that are due to the high correlation between genetic markers, as well as decrease computational burden, and relax the excessive multiple testing correction. LD pruning was done as described in Otyama et al. (2020) using LD r^2^ threshold of 0.2. Tests were carried out in PLINK version 1.9 (Purcell et al. 2007) without controlling for confounders and using the Genome-Wide Interaction Analysis Software, CASSI, controlling for population stratification using PCA (CASSI ; Ueki and Cordell 2012). For each trait we fit a linear regression model:

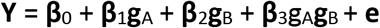

In this model, for each inspected SNP pair (**A**, **B**), **g**_A_ and **g**_B_ represent allele counts; **β**_0_ is the overall mean; **β**_1_ and **β**_2_ are the additive effects of SNPs A and B; **β**_3_ is the interaction coefficient tested for significance; and **e** is the random error following N(0, **σ**^2^_e_). To correct for multiple comparisons, a Bonferroni threshold (α = 0.05) was applied to all p-values and only interactions with p-value < 4.65E-08 were considered significantly associated with a trait.

### Enrichment Analysis

For each trait, P values < 1.54E-05 from GWAS and P values < 4.65E-08 from GWASi were used to identify significant SNPs and SNP pair interactions. All genes within a 1000 kbp window around GWAS SNPs and SNP pair interactions were extracted. Enrichment analysis and the assignment of genes to functional categories was implemented in PeanutMine and BLUEGENES from InterMine (Smith et al. 2012), on PeanutBase (Berendzen et al. 2021; Dash et al. 2016).

Significantly enriched gene ontology (GO) terms, gene families, and pathways in the gene list compared with background genes were defined with a hypergeometric test at p < 0.05.

Background genes were defined as all annotated genes in Tifrunner genome 1 annotation 2 (Dash et al. 2016). The resulting P values from enrichment analyses were corrected for type I error due to multiple testing using the Benjamin Hochberg method.

## Results and Discussion

### Natural variation of fatty acids in peanut seed oil

The relative concentration of eight fatty acid components (palmitic [C16:0], stearic [C18:0], oleic [C18:1], linoleic [C18:2), arachidic [C20:0], gadoleic [C20:1], behenic [C22:0], and lignoceric [C24:0]), which are the most commonly found fatty acids in peanut, were determined from oil extracted from dried, harvested seeds (Dezern 2018; Otyama et al. 2019). The total unsaturated fatty acid content was calculated as the sum of oleic, linoleic, and gadoleic concentrations and total saturated fatty acid content was calculated as the sum of palmitic, stearic, lignoceric, behenic, and arachidic fatty acid concentrations observed in the oil.

We observed significant differences between the performances of replicated genotypes in each environment for all measured traits at p < 0.01 (**Table 1**), suggesting a significant gene by environment interaction (GxE) effect on fatty acid content. Trait means ranged from 56.17 for oleic acid content all the way down to 1.27 for lignoceric content. Oleic and linoleic acid content had the highest heritability estimates (0.95) followed by palmitic (0.94), stearic (0.86), and gadoleic acid content (0.85). Lignoceric and arachidic acids had the lowest heritability estimates at 0.48 and 0.33, respectively (**Table 1**). This indicates high repeatability of the measured traits for the primary fatty acids detected in peanuts and suggests a strong influence of the genetics on the observed natural variation in fatty acid composition.

**Table 1.** Natural variation of fatty acid components in peanut seed oil.

| Trait | Heritability (H<sup>2</sup>) | Genotype Variance | Residual Variance | Grand Mean | LSD | CV | Genotype significance |  |
| --- | --- | --- | --- | --- | --- | --- | --- | --- |
| Oleic | 0.95 | 81.53 | 12.42 | 56.17 | 5.46 | 6.27 | 0.000 | ** |
| Linoleic | 0.95 | 54.89 | 8.28 | 24.37 | 4.46 | 11.81 | 0.000 | ** |
| Gadoleic | 0.85 | 0.04 | 0.02 | 0.99 | 0.22 | 15.28 | 0.000 | ** |
| Palmitic | 0.94 | 2.22 | 0.41 | 9.87 | 0.99 | 6.51 | 0.000 | ** |
| Stearic | 0.86 | 0.40 | 0.20 | 3.15 | 0.66 | 14.18 | 0.000 | ** |
| Arachidic | 0.33 | 0.06 | 0.35 | 1.49 | 0.54 | 39.44 | 0.008 | ** |
| Behenic | 0.69 | 0.11 | 0.16 | 2.67 | 0.52 | 14.80 | 0.000 | ** |
| Lignoceric | 0.48 | 0.02 | 0.06 | 1.27 | 0.28 | 19.95 | 0.000 | ** |

n = 3 replicates; H<sup>2</sup> = broad-sense heritability; LSD = least significant difference at 5%; CV = coefficient of variation as a percentage where $CV = ((VMSE)/Grand\ Mean) * 100$ . $LSD = t(1-0.05, df_{Err}) \times ASED$ , where ASED = average standard error of the differences of the means and $df_{Err}$ = error degrees of freedom. \*\* Significant genotype difference at $\alpha = 0.001$ .

Pearson correlation coefficients were calculated between traits to give a measure of the strength of linear associations. Color intensity and the size of the circle are proportional to the correlation coefficients at significance level α = 0.01 (**Figure S1. A**). We detected a strong negative correlation between oleic acid and linoleic acid (r^2^ = 0.91) and between oleic acid and palmitic acid (r^2^ = 0.67) **(Figure S1. A, Table S1**). This is consistent with earlier studies (Barkley et al. 2011; Barkley et al. 2013; Isleib et al. 2006; Nawade et al. 2016; Wang et al. 2015). The content of oleic and palmitic acid strongly influenced the total unsaturated and saturated fatty acid content of the seed oil, respectively, due to their relative abundance in peanut seed oil. Gadoleic content was positively correlated with behenic acid content (r^2^ = 0.56), but showed no significant correlation with lignoceric acid. We also observed a moderately low positive correlation between linoleic and palmitic acid content (r^2^ = 0.47) and a moderately high positive correlation between arachidic and behenic acid content (r^2^ = 0.63) (**Figure S1. A, Table S1**). These results imply a shared genetic architecture among traits showing high correlations which we sought to investigate further in the subsequent subsections.

### LD and Population structure

In the 885 unique genotypes, linkage disequilibrium (measured as r^2^), decayed to a baseline of 0.2 at 4.8 Mbp (**Figure S1. B**). This is relatively high compared to other self-pollinating species, and is likely strongly influenced by peanut’s recent tetraploid history (Bertioli et al. 2016; Bertioli et al. 2019). We previously reported the genetic diversity in the USDA peanut core collection is best explained by four major clades, generally corresponding with peanut subspecies classification and botanical varieties from which four main market types – Runner, Virginia, Valencia, and Spanish – are defined (Otyama et al. 2020; Otyama et al. 2019).

### Genomic regions and variants associated with fatty acid concentration

GWAS was conducted for each trait using the Fixed and random model Circulating Probability Unification model (FarmCPU) fitting principal components from PCA to control for stratification in the data.

GWAS was both sensitive to population structure and population size based on QQ plots, judged from fitting models with varying PCs ranging 0 to 4 and population size representing 25%, 50%, and 75% of all genotyped and phenotyped individuals (**Table S1** – underlying data not presented).

We detected 251 SNPs with significant trait associations with different fatty acid components (**α** = 5% for P values < 1.54E-05 and FDR adjusted P values < 1.0E-02). Of the significant associations, 12 SNPs affected the variation of joint trait pairs while two other SNPs affected trait trios. The trait pairs affected by pleiotropic SNPs include: oleic-linoleic acid, saturated-unsaturated fatty acids, unsaturated fatty acid-linoleic acid, and unsaturated fatty acid-palmitic acid content. The trait trios affected jointly include: oleic-linoleic-palmitic acid, and behenic- gadoleic-lignoceric acid content (**Table 2**, **Figure 5**). The remaining 237 SNPs were significantly associated with individual fatty acid components (**Table S2**).

**Table 2:**
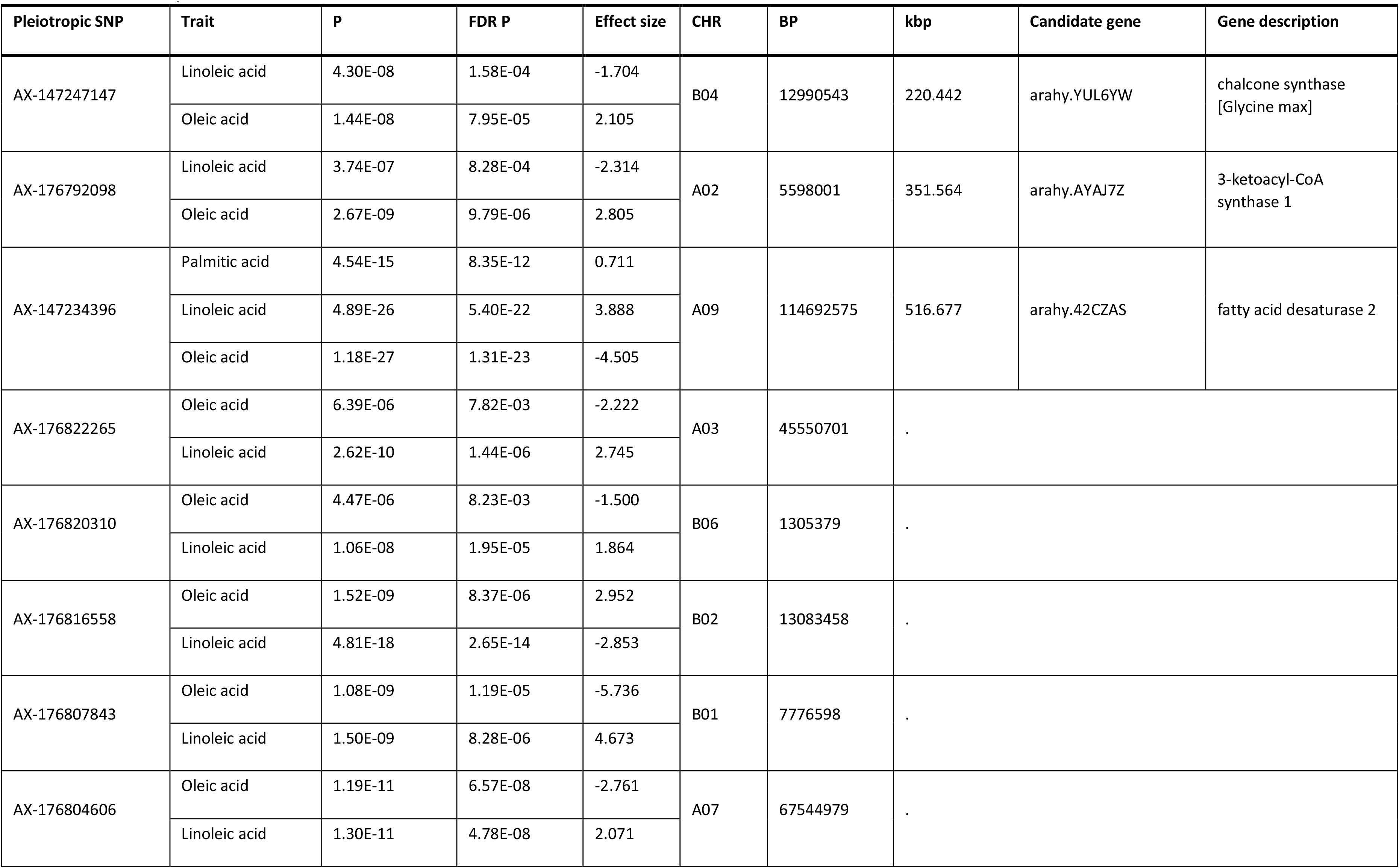
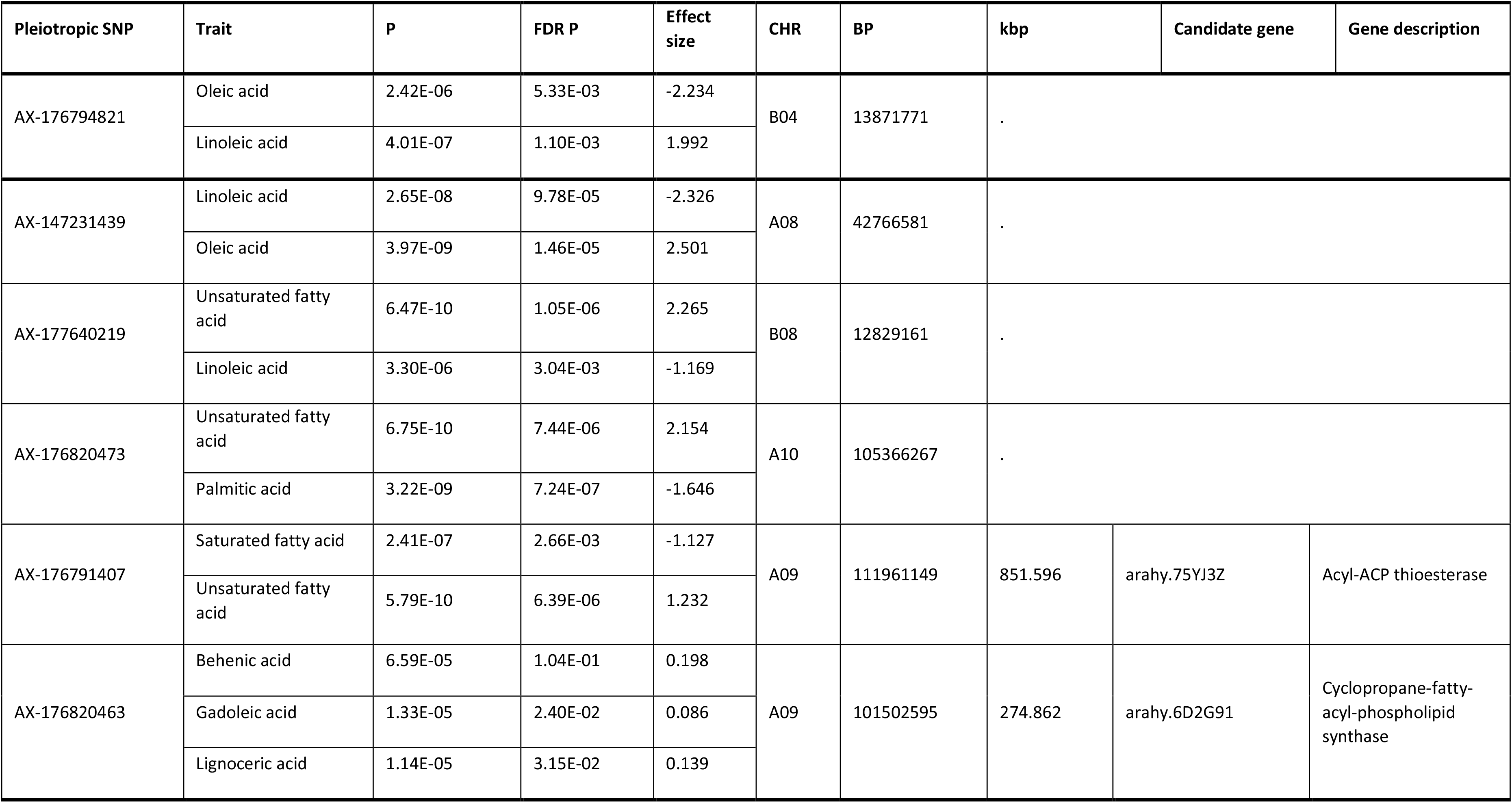
Pleiotropic variants and traits.

| Pleiotropic SNP | Trait | P | FDR P | Effect size | CHR | BP | kbp | Candidate gene | Gene description |
| --- | --- | --- | --- | --- | --- | --- | --- | --- | --- |
| AX-147247147 | Linoleic acid | 4.30E-08 | 1.58E-04 | -1.704 | B04 | 12990543 | 220.442 | arahy.YUL6YW | chalcone synthase<br>[Glycine max] |
|  | Oleic acid | 1.44E-08 | 7.95E-05 | 2.105 |  |  |  |  |  |
| AX-176792098 | Linoleic acid | 3.74E-07 | 8.28E-04 | -2.314 | A02 | 5598001 | 351.564 | arahy.AYAJ7Z | 3-ketoacyl-CoA<br>synthase 1 |
|  | Oleic acid | 2.67E-09 | 9.79E-06 | 2.805 |  |  |  |  |  |
| AX-147234396 | Palmitic acid | 4.54E-15 | 8.35E-12 | 0.711 | A09 | 114692575 | 516.677 | arahy.42CZAS | fatty acid desaturase 2 |
|  | Linoleic acid | 4.89E-26 | 5.40E-22 | 3.888 |  |  |  |  |  |
|  | Oleic acid | 1.18E-27 | 1.31E-23 | -4.505 |  |  |  |  |  |
| AX-176822265 | Oleic acid | 6.39E-06 | 7.82E-03 | -2.222 | A03 | 45550701 | . |  |  |
|  | Linoleic acid | 2.62E-10 | 1.44E-06 | 2.745 |  |  |  |  |  |
| AX-176820310 | Oleic acid | 4.47E-06 | 8.23E-03 | -1.500 | B06 | 1305379 | . |  |  |
|  | Linoleic acid | 1.06E-08 | 1.95E-05 | 1.864 |  |  |  |  |  |
| AX-176816558 | Oleic acid | 1.52E-09 | 8.37E-06 | 2.952 | B02 | 13083458 | . |  |  |
|  | Linoleic acid | 4.81E-18 | 2.65E-14 | -2.853 |  |  |  |  |  |
| AX-176807843 | Oleic acid | 1.08E-09 | 1.19E-05 | -5.736 | B01 | 7776598 | . |  |  |
|  | Linoleic acid | 1.50E-09 | 8.28E-06 | 4.673 |  |  |  |  |  |
| AX-176804606 | Oleic acid | 1.19E-11 | 6.57E-08 | -2.761 | A07 | 67544979 | . |  |  |
|  | Linoleic acid | 1.30E-11 | 4.78E-08 | 2.071 |  |  |  |  |  |
| AX-176794821 | Oleic acid | 2.42E-06 | 5.33E-03 | -2.234 | B04 | 13871771 | . |  |  |
|  | Linoleic acid | 4.01E-07 | 1.10E-03 | 1.992 |  |  |  |  |  |
| AX-147231439 | Linoleic acid | 2.65E-08 | 9.78E-05 | -2.326 | A08 | 42766581 | . |  |  |
|  | Oleic acid | 3.97E-09 | 1.46E-05 | 2.501 |  |  |  |  |  |
| AX-177640219 | Unsaturated fatty acid | 6.47E-10 | 1.05E-06 | 2.265 | B08 | 12829161 | . |  |  |
|  | Linoleic acid | 3.30E-06 | 3.04E-03 | -1.169 |  |  |  |  |  |
| AX-176820473 | Unsaturated fatty acid | 6.75E-10 | 7.44E-06 | 2.154 | A10 | 105366267 | . |  |  |
|  | Palmitic acid | 3.22E-09 | 7.24E-07 | -1.646 |  |  |  |  |  |
| AX-176791407 | Saturated fatty acid | 2.41E-07 | 2.66E-03 | -1.127 | A09 | 111961149 | 851.596 | arahy.75YJ3Z | Acyl-ACP thioesterase |
|  | Unsaturated fatty acid | 5.79E-10 | 6.39E-06 | 1.232 |  |  |  |  |  |
| AX-176820463 | Behenic acid | 6.59E-05 | 1.04E-01 | 0.198 | A09 | 101502595 | 274.862 | arahy.6D2G91 | Cyclopropane-fatty-acyl-phospholipid synthase |
|  | Gadoleic acid | 1.33E-05 | 2.40E-02 | 0.086 |  |  |  |  |  |
|  | Lignoceric acid | 1.14E-05 | 3.15E-02 | 0.139 |  |  |  |  |  |

In 10 of the 14 pleiotropic cases detected, the SNPs are jointly associated with oleic and linoleic acid with contrasting effects - a SNP associated with increased oleic acid content also associated with decreased linoleic acid (**Table 2**, **Figure 5. A-J**). Significant SNPs were distributed throughout the genome. We further evaluated SNPs which were located within 1000 kbp of 55 genes with predicted fatty acid biosynthesis function (**Table 3**).

**Table 3.**
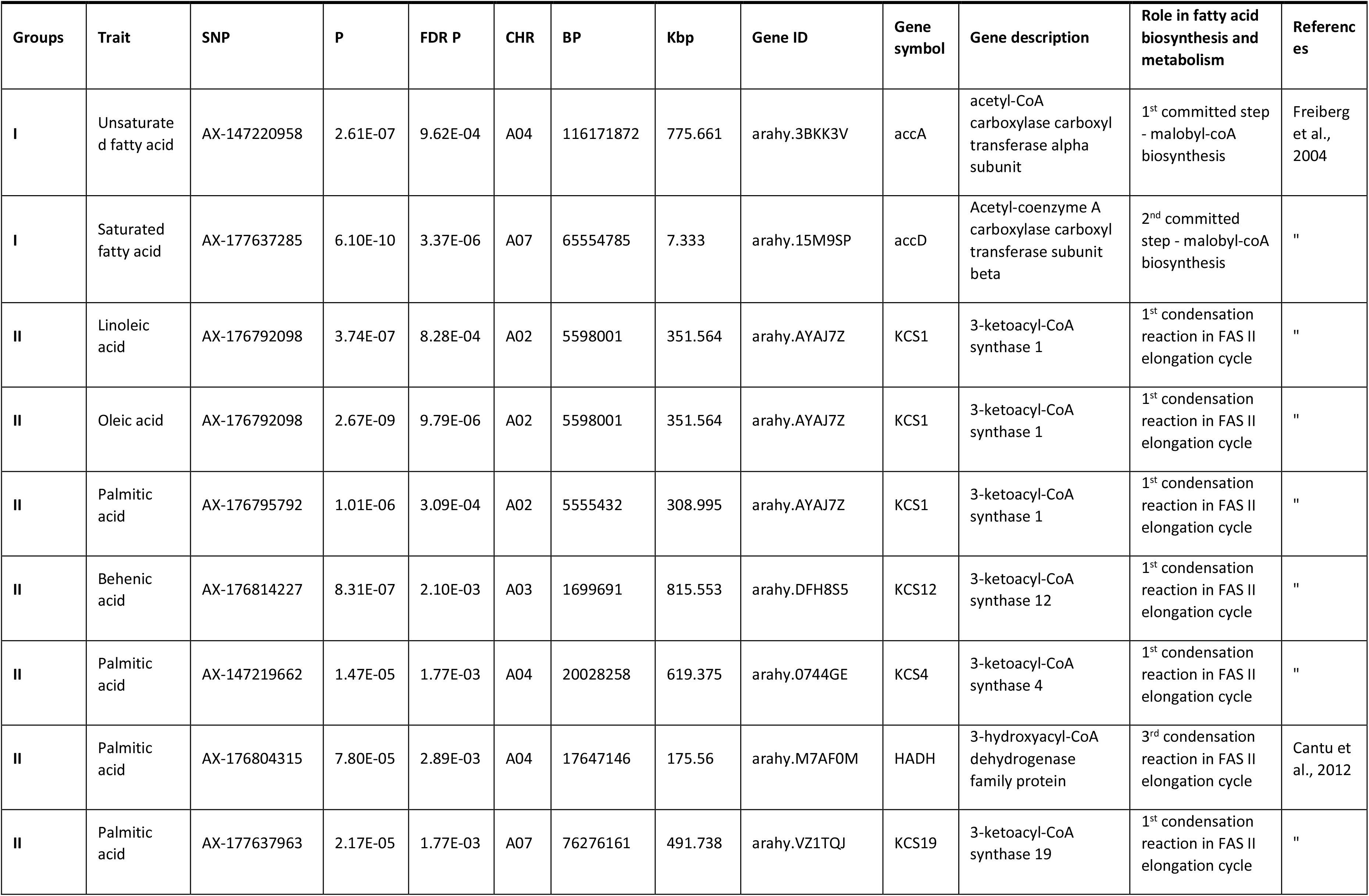
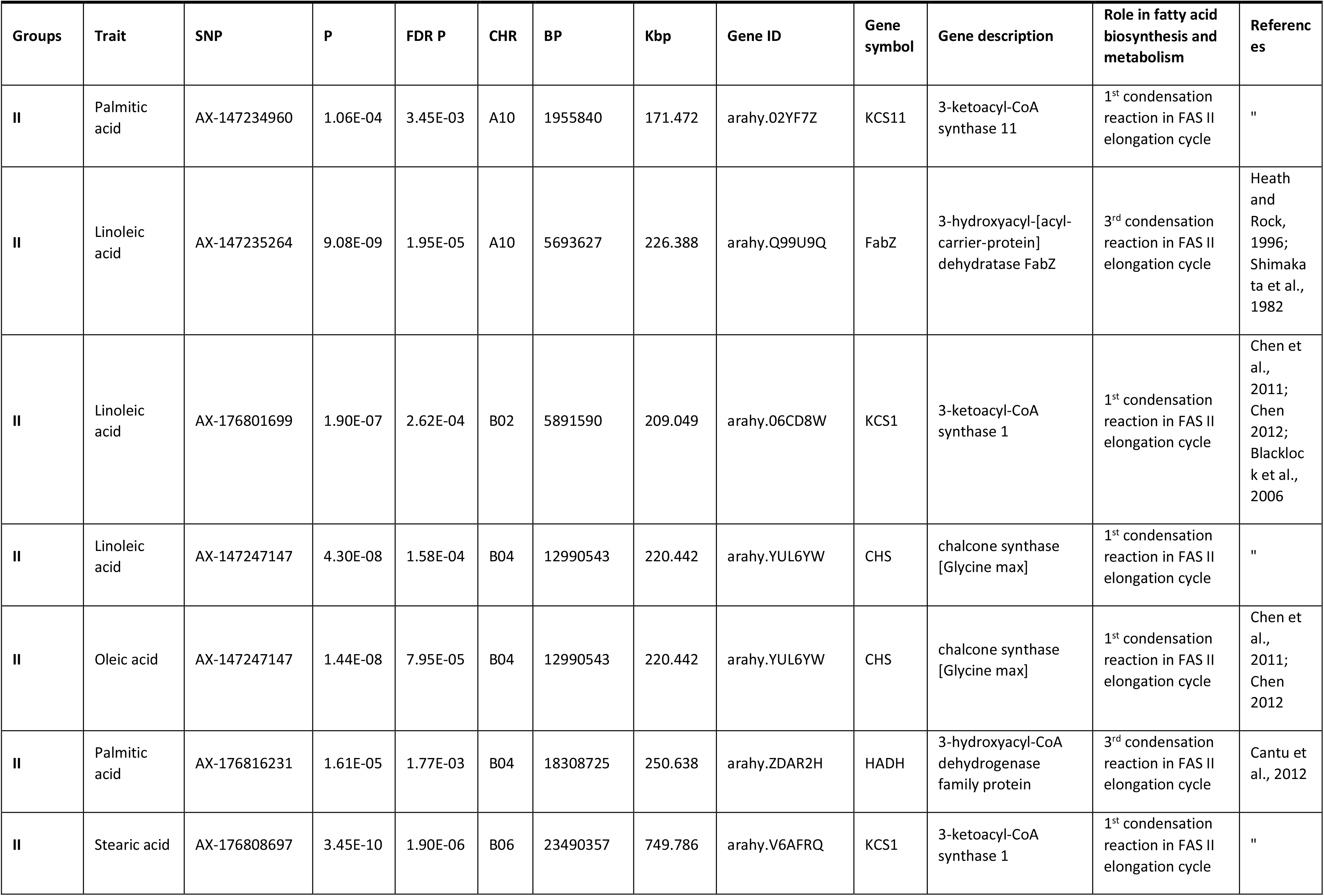
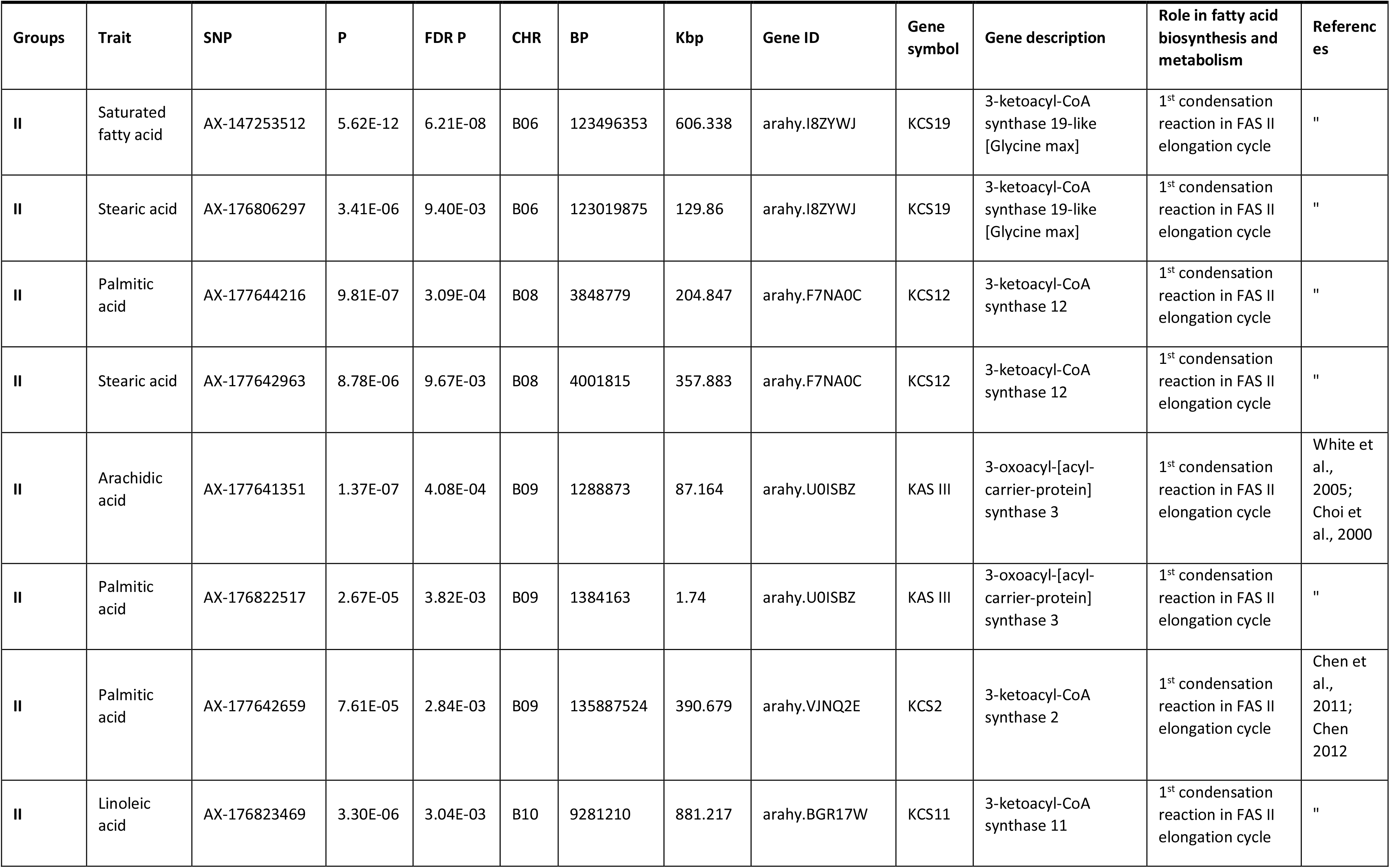
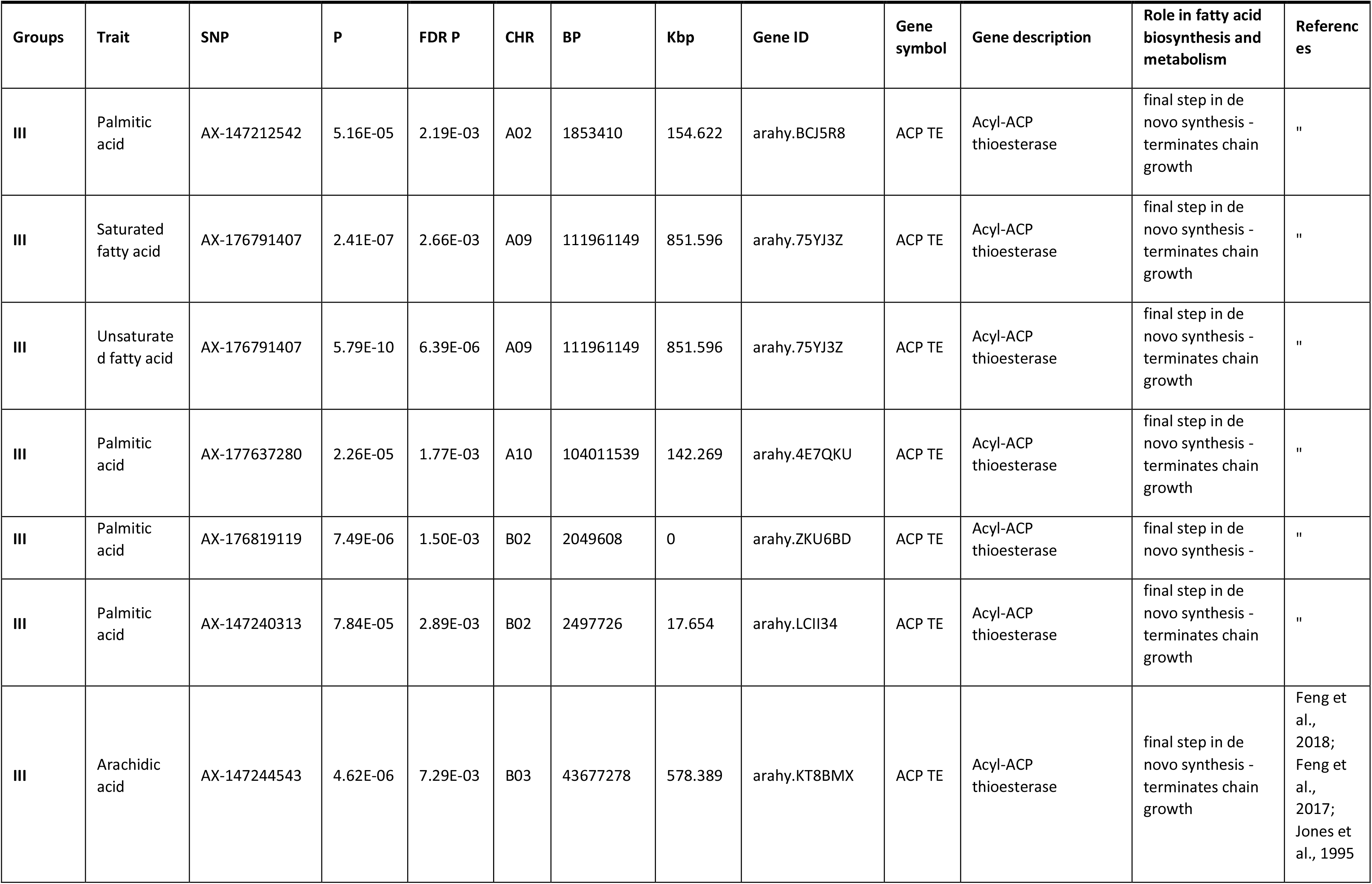
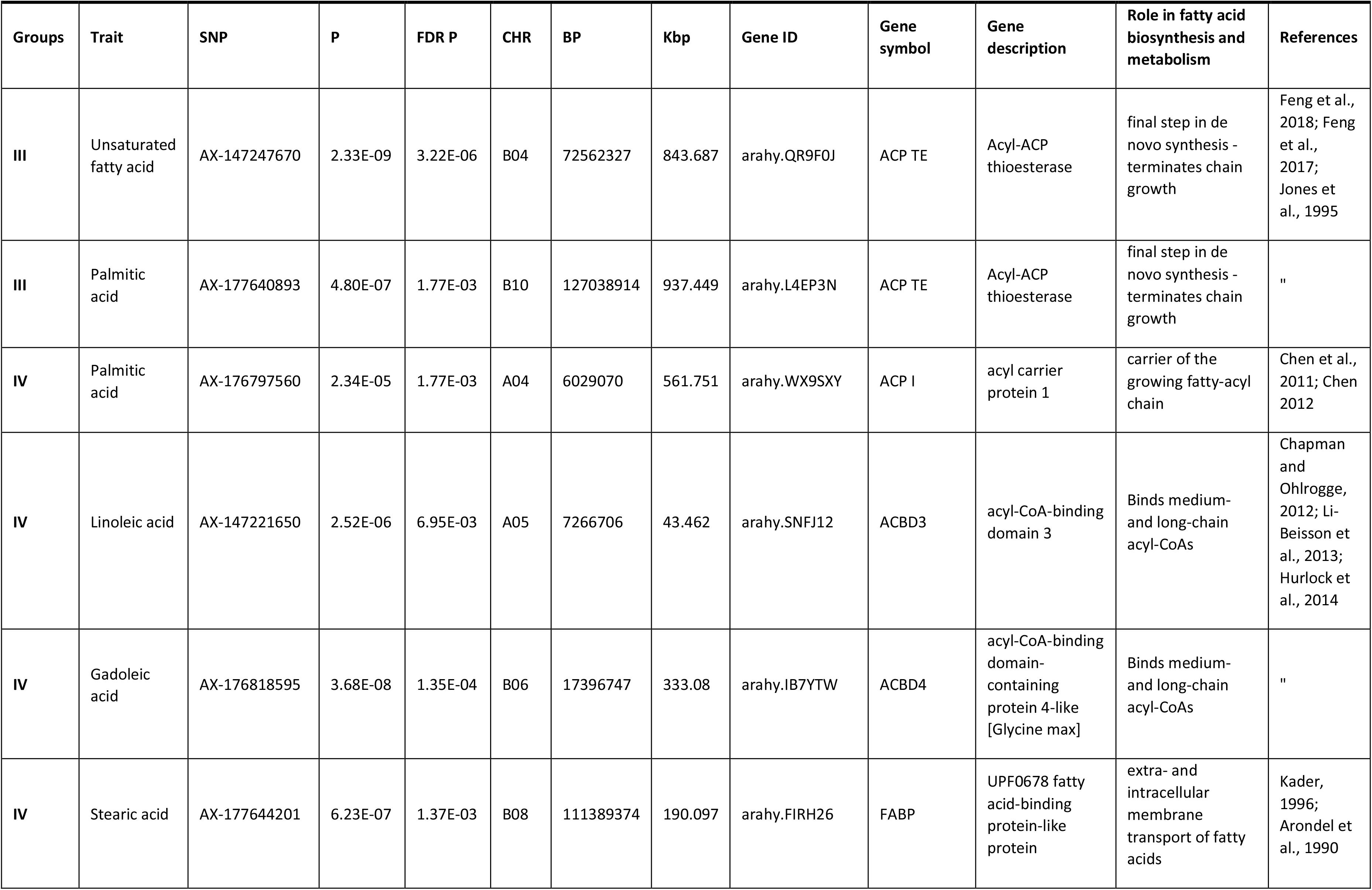
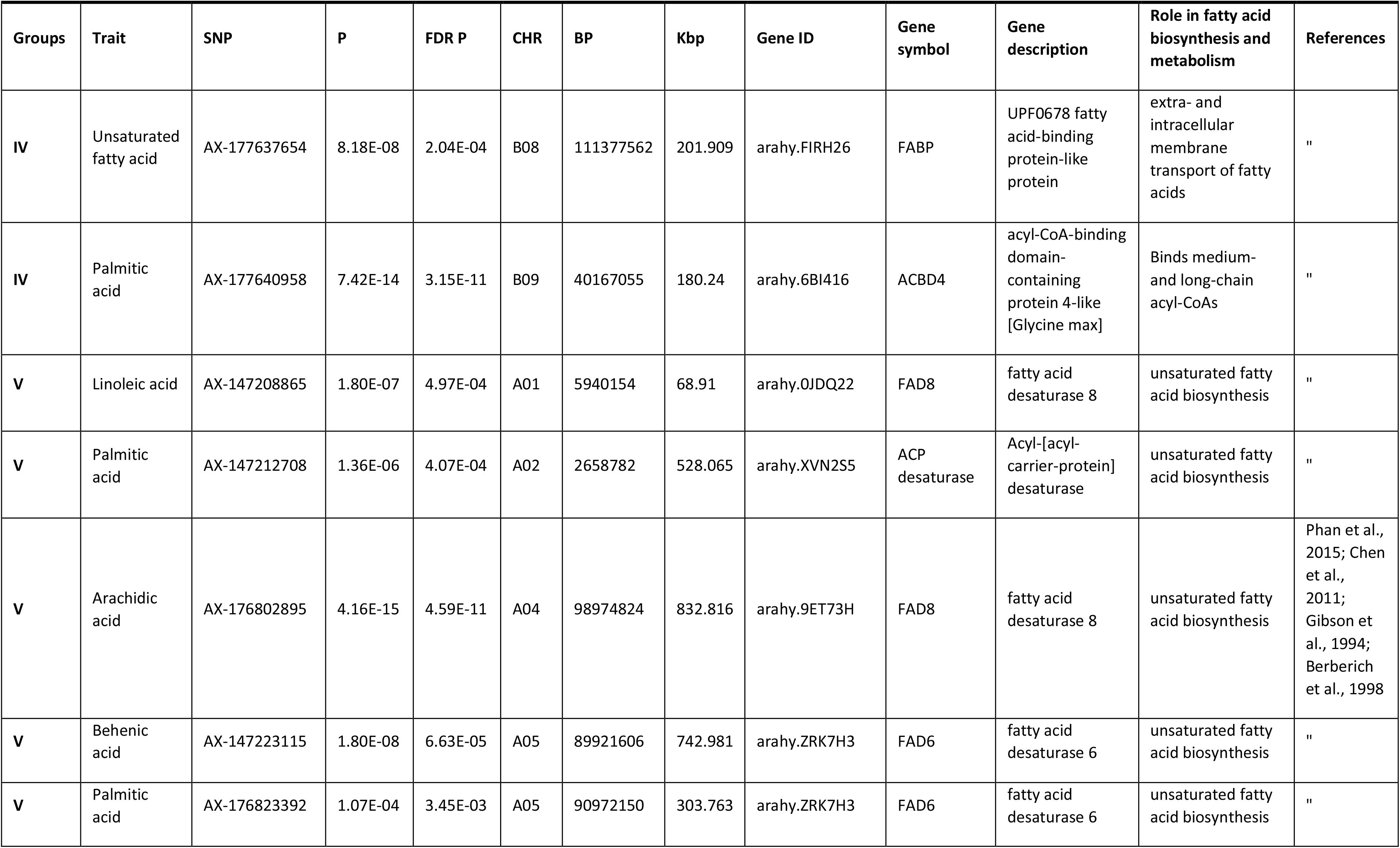
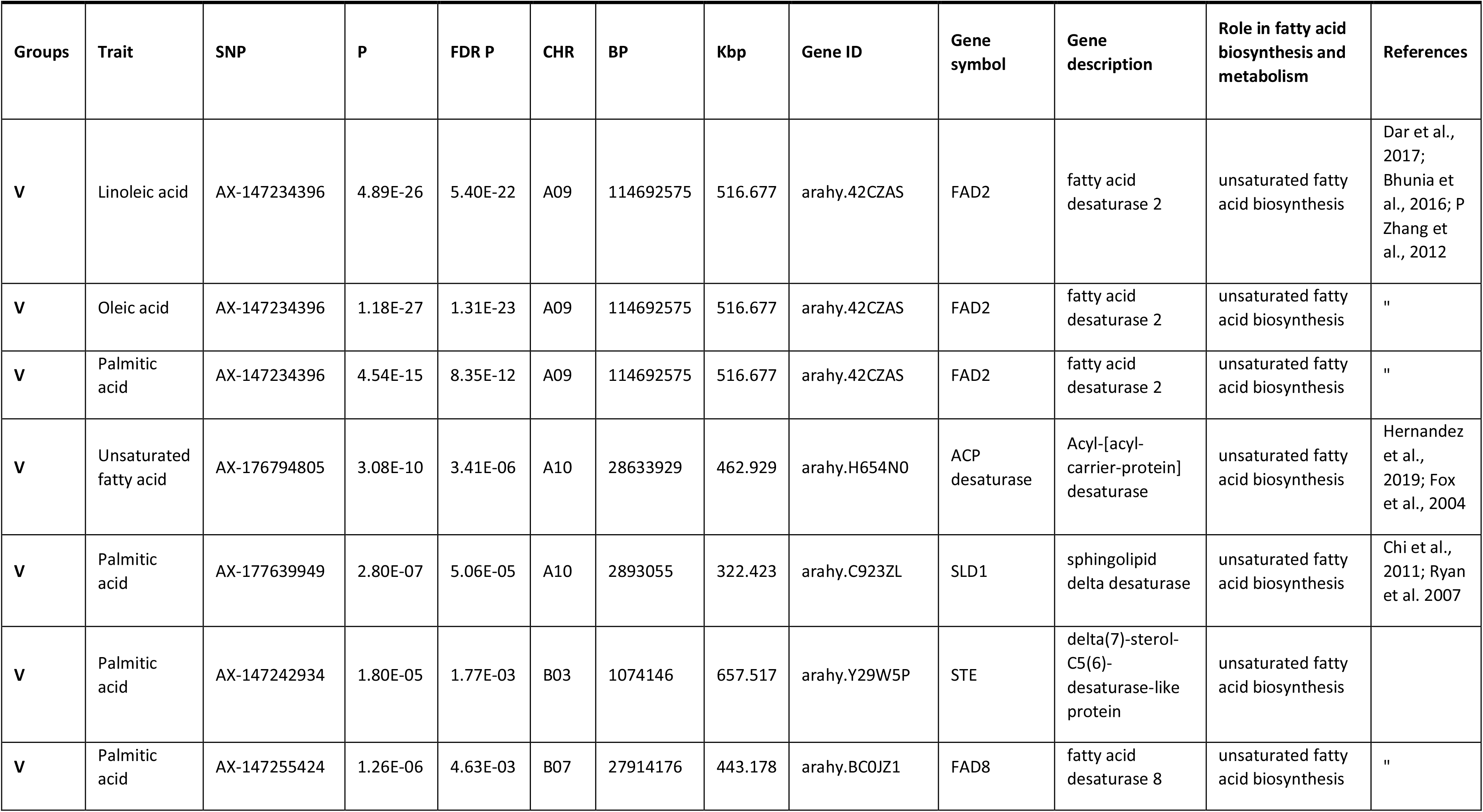
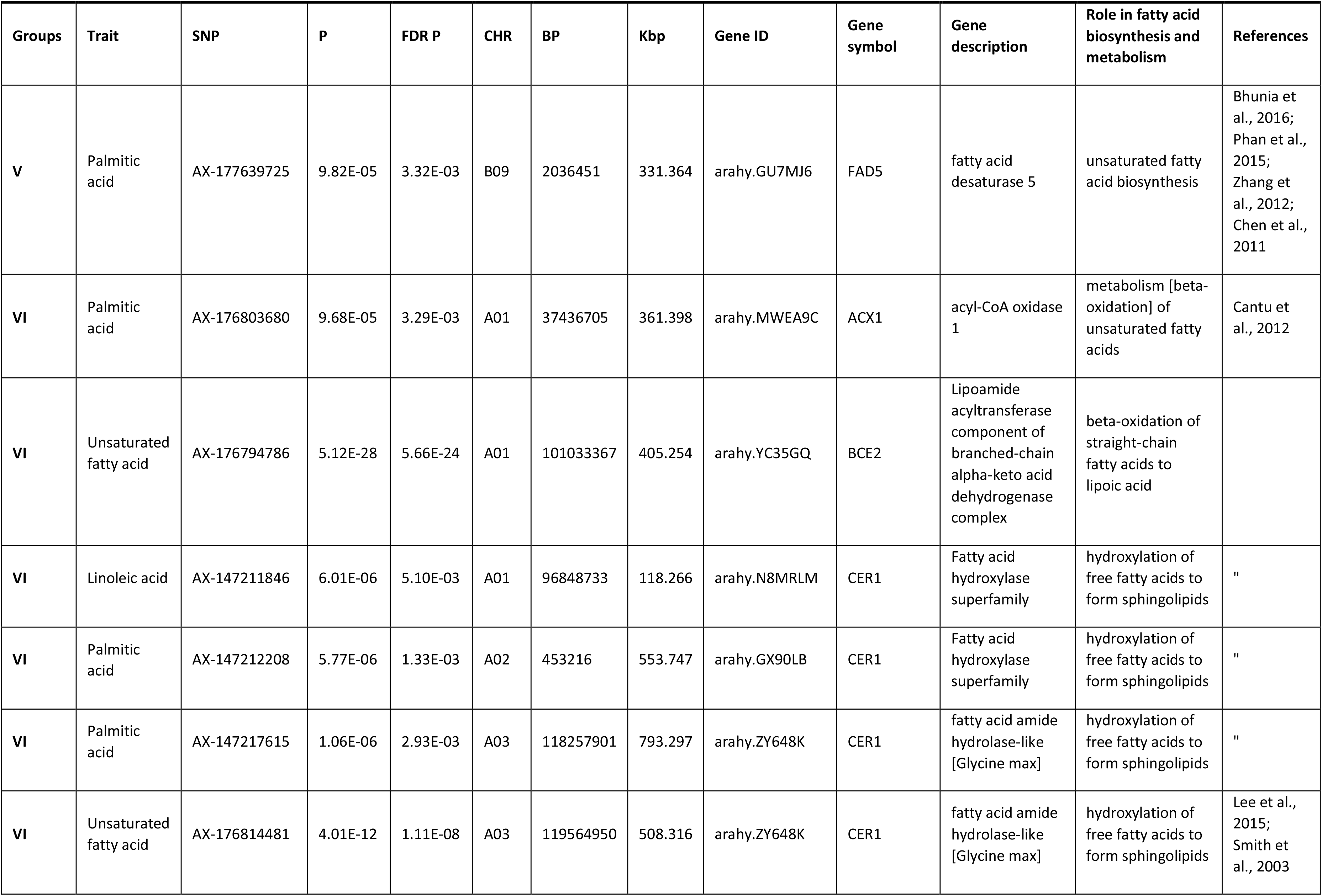
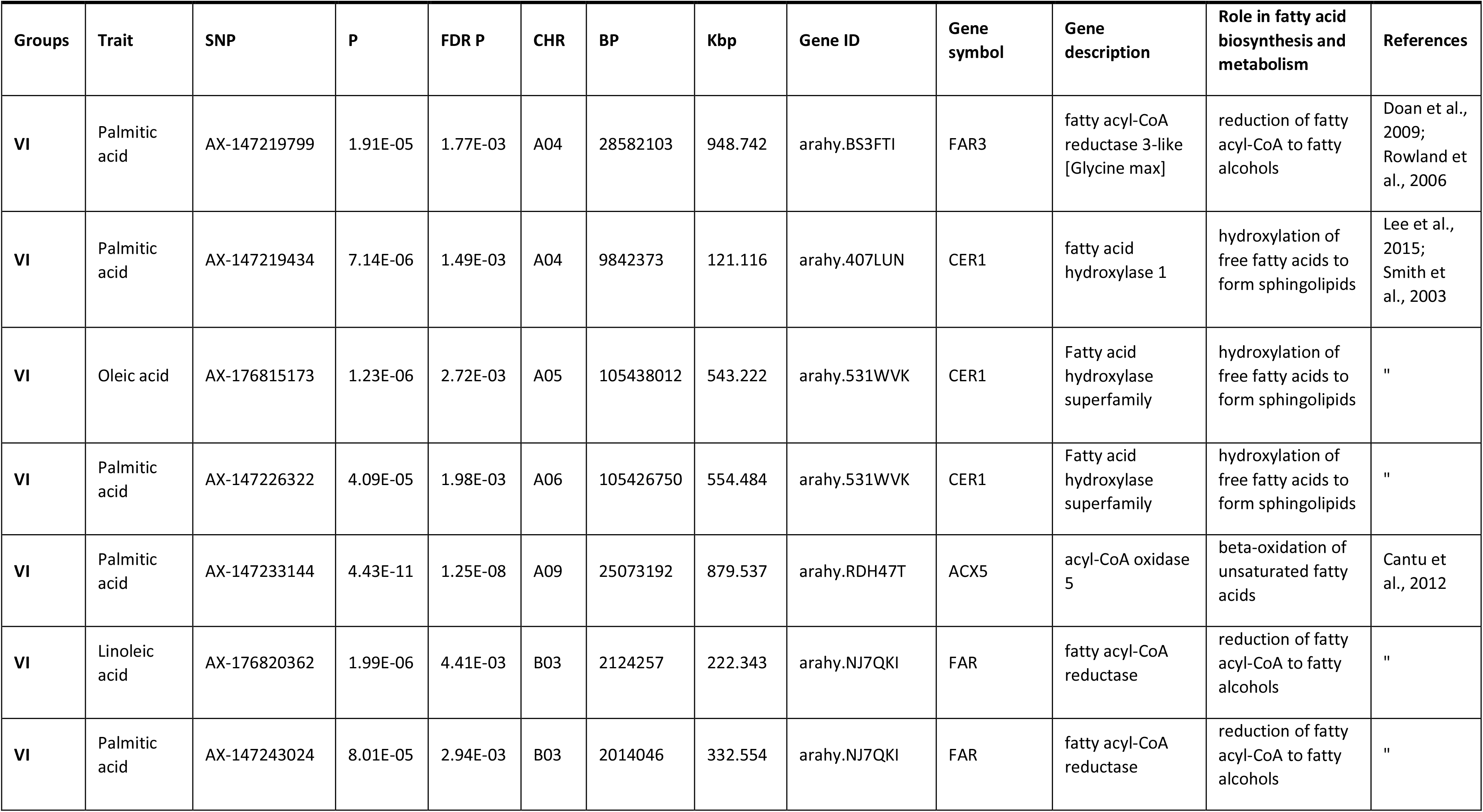
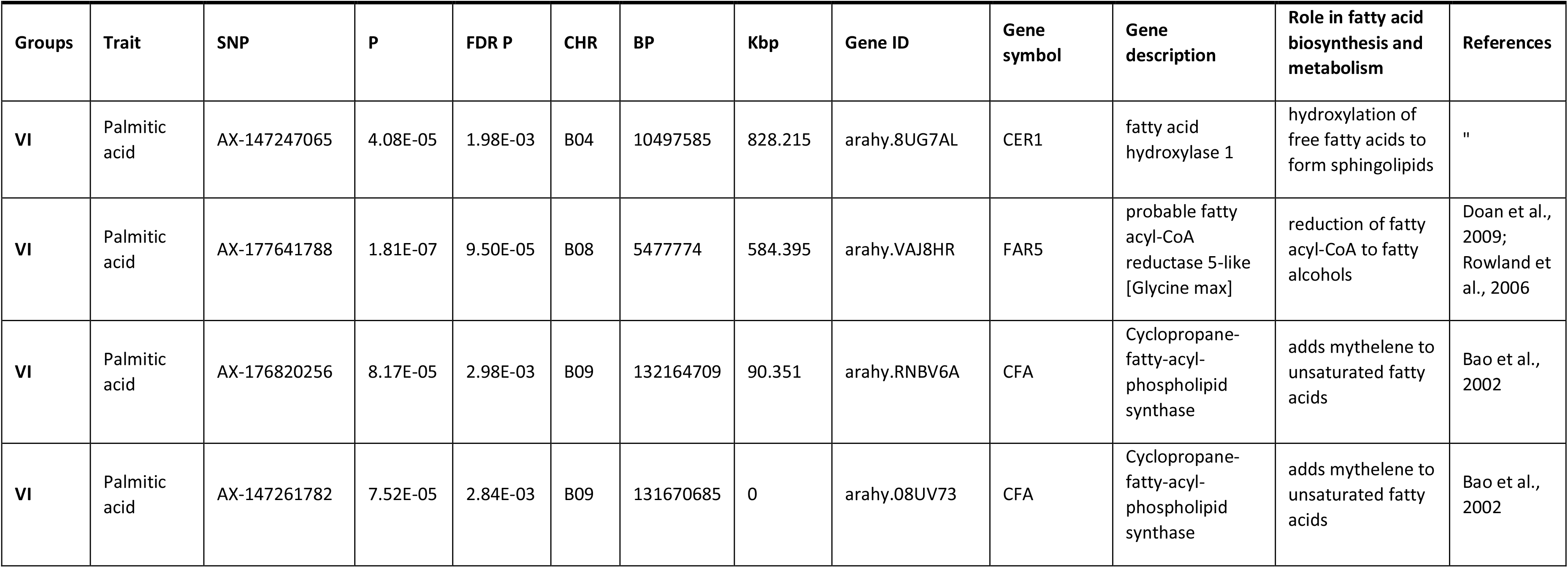
GWAS SNPs within 1000 kbp of known fatty acid biosynthesis and fatty acid metabolism genes.

| Groups | Trait | SNP | P | FDR P | CHR | BP | Kbp | Gene ID | Gene symbol | Gene description | Role in fatty acid biosynthesis and metabolism | References |
| --- | --- | --- | --- | --- | --- | --- | --- | --- | --- | --- | --- | --- |
| I | Unsaturated fatty acid | AX-147220958 | 2.61E-07 | 9.62E-04 | A04 | 116171872 | 775.661 | arahy.3BKK3V | accA | acetyl-CoA carboxylase carboxyl transferase alpha subunit | 1<sup>st</sup> committed step - malonyl-CoA biosynthesis | Freiberg et al., 2004 |
| I | Saturated fatty acid | AX-177637285 | 6.10E-10 | 3.37E-06 | A07 | 65554785 | 7.333 | arahy.15M9SP | accD | Acetyl-coenzyme A carboxylase carboxyl transferase subunit beta | 2<sup>nd</sup> committed step - malonyl-CoA biosynthesis | " |
| II | Linoleic acid | AX-176792098 | 3.74E-07 | 8.28E-04 | A02 | 5598001 | 351.564 | arahy.AYAJ7Z | KCS1 | 3-ketoacyl-CoA synthase 1 | 1<sup>st</sup> condensation reaction in FAS II elongation cycle | " |
| II | Oleic acid | AX-176792098 | 2.67E-09 | 9.79E-06 | A02 | 5598001 | 351.564 | arahy.AYAJ7Z | KCS1 | 3-ketoacyl-CoA synthase 1 | 1<sup>st</sup> condensation reaction in FAS II elongation cycle | " |
| II | Palmitic acid | AX-176795792 | 1.01E-06 | 3.09E-04 | A02 | 5555432 | 308.995 | arahy.AYAJ7Z | KCS1 | 3-ketoacyl-CoA synthase 1 | 1<sup>st</sup> condensation reaction in FAS II elongation cycle | " |
| II | Behenic acid | AX-176814227 | 8.31E-07 | 2.10E-03 | A03 | 1699691 | 815.553 | arahy.DFH8S5 | KCS12 | 3-ketoacyl-CoA synthase 12 | 1<sup>st</sup> condensation reaction in FAS II elongation cycle | " |
| II | Palmitic acid | AX-147219662 | 1.47E-05 | 1.77E-03 | A04 | 20028258 | 619.375 | arahy.0744GE | KCS4 | 3-ketoacyl-CoA synthase 4 | 1<sup>st</sup> condensation reaction in FAS II elongation cycle | " |
| II | Palmitic acid | AX-176804315 | 7.80E-05 | 2.89E-03 | A04 | 17647146 | 175.56 | arahy.M7AF0M | HADH | 3-hydroxyacyl-CoA dehydrogenase family protein | 3<sup>rd</sup> condensation reaction in FAS II elongation cycle | Cantu et al., 2012 |
| II | Palmitic acid | AX-177637963 | 2.17E-05 | 1.77E-03 | A07 | 76276161 | 491.738 | arahy.VZ1TQJ | KCS19 | 3-ketoacyl-CoA synthase 19 | 1<sup>st</sup> condensation reaction in FAS II elongation cycle | " |
| II | Palmitic acid | AX-147234960 | 1.06E-04 | 3.45E-03 | A10 | 1955840 | 171.472 | arahy.02YF7Z | KCS11 | 3-ketoacyl-CoA synthase 11 | 1<sup>st</sup> condensation reaction in FAS II elongation cycle | " |
| II | Linoleic acid | AX-147235264 | 9.08E-09 | 1.95E-05 | A10 | 5693627 | 226.388 | arahy.Q99U9Q | FabZ | 3-hydroxyacyl-[acyl-carrier-protein] dehydratase FabZ | 3<sup>rd</sup> condensation reaction in FAS II elongation cycle | Heath and Rock, 1996; Shimakata et al., 1982 |
| II | Linoleic acid | AX-176801699 | 1.90E-07 | 2.62E-04 | B02 | 5891590 | 209.049 | arahy.06CD8W | KCS1 | 3-ketoacyl-CoA synthase 1 | 1<sup>st</sup> condensation reaction in FAS II elongation cycle | Chen et al., 2011; Chen 2012; Blacklock et al., 2006 |
| II | Linoleic acid | AX-147247147 | 4.30E-08 | 1.58E-04 | B04 | 12990543 | 220.442 | arahy.YUL6YW | CHS | chalcone synthase [Glycine max] | 1<sup>st</sup> condensation reaction in FAS II elongation cycle | " |
| II | Oleic acid | AX-147247147 | 1.44E-08 | 7.95E-05 | B04 | 12990543 | 220.442 | arahy.YUL6YW | CHS | chalcone synthase [Glycine max] | 1<sup>st</sup> condensation reaction in FAS II elongation cycle | Chen et al., 2011; Chen 2012 |
| II | Palmitic acid | AX-176816231 | 1.61E-05 | 1.77E-03 | B04 | 18308725 | 250.638 | arahy.ZDAR2H | HADH | 3-hydroxyacyl-CoA dehydrogenase family protein | 3<sup>rd</sup> condensation reaction in FAS II elongation cycle | Cantu et al., 2012 |
| II | Stearic acid | AX-176808697 | 3.45E-10 | 1.90E-06 | B06 | 23490357 | 749.786 | arahy.V6AFRQ | KCS1 | 3-ketoacyl-CoA synthase 1 | 1<sup>st</sup> condensation reaction in FAS II elongation cycle | " |
| II | Saturated fatty acid | AX-147253512 | 5.62E-12 | 6.21E-08 | B06 | 123496353 | 606.338 | arahy.I8ZYWJ | KCS19 | 3-ketoacyl-CoA synthase 19-like [Glycine max] | 1<sup>st</sup> condensation reaction in FAS II elongation cycle | " |
| II | Stearic acid | AX-176806297 | 3.41E-06 | 9.40E-03 | B06 | 123019875 | 129.86 | arahy.I8ZYWJ | KCS19 | 3-ketoacyl-CoA synthase 19-like [Glycine max] | 1<sup>st</sup> condensation reaction in FAS II elongation cycle | " |
| II | Palmitic acid | AX-177644216 | 9.81E-07 | 3.09E-04 | B08 | 3848779 | 204.847 | arahy.F7NAOC | KCS12 | 3-ketoacyl-CoA synthase 12 | 1<sup>st</sup> condensation reaction in FAS II elongation cycle | " |
| II | Stearic acid | AX-177642963 | 8.78E-06 | 9.67E-03 | B08 | 4001815 | 357.883 | arahy.F7NAOC | KCS12 | 3-ketoacyl-CoA synthase 12 | 1<sup>st</sup> condensation reaction in FAS II elongation cycle | " |
| II | Arachidic acid | AX-177641351 | 1.37E-07 | 4.08E-04 | B09 | 1288873 | 87.164 | arahy.U0ISBZ | KAS III | 3-oxoacyl-[acyl-carrier-protein] synthase 3 | 1<sup>st</sup> condensation reaction in FAS II elongation cycle | White et al., 2005; Choi et al., 2000 |
| II | Palmitic acid | AX-176822517 | 2.67E-05 | 3.82E-03 | B09 | 1384163 | 1.74 | arahy.U0ISBZ | KAS III | 3-oxoacyl-[acyl-carrier-protein] synthase 3 | 1<sup>st</sup> condensation reaction in FAS II elongation cycle | " |
| II | Palmitic acid | AX-177642659 | 7.61E-05 | 2.84E-03 | B09 | 135887524 | 390.679 | arahy.VJNQ2E | KCS2 | 3-ketoacyl-CoA synthase 2 | 1<sup>st</sup> condensation reaction in FAS II elongation cycle | Chen et al., 2011; Chen 2012 |
| II | Linoleic acid | AX-176823469 | 3.30E-06 | 3.04E-03 | B10 | 9281210 | 881.217 | arahy.BGR17W | KCS11 | 3-ketoacyl-CoA synthase 11 | 1<sup>st</sup> condensation reaction in FAS II elongation cycle | " |
| III | Palmitic acid | AX-147212542 | 5.16E-05 | 2.19E-03 | A02 | 1853410 | 154.622 | arahy.BCJ5R8 | ACP TE | Acyl-ACP thioesterase | final step in de novo synthesis - terminates chain growth | " |
| III | Saturated fatty acid | AX-176791407 | 2.41E-07 | 2.66E-03 | A09 | 111961149 | 851.596 | arahy.75YJ3Z | ACP TE | Acyl-ACP thioesterase | final step in de novo synthesis - terminates chain growth | " |
| III | Unsaturated fatty acid | AX-176791407 | 5.79E-10 | 6.39E-06 | A09 | 111961149 | 851.596 | arahy.75YJ3Z | ACP TE | Acyl-ACP thioesterase | final step in de novo synthesis - terminates chain growth | " |
| III | Palmitic acid | AX-177637280 | 2.26E-05 | 1.77E-03 | A10 | 104011539 | 142.269 | arahy.4E7QKU | ACP TE | Acyl-ACP thioesterase | final step in de novo synthesis - terminates chain growth | " |
| III | Palmitic acid | AX-176819119 | 7.49E-06 | 1.50E-03 | B02 | 2049608 | 0 | arahy.ZKU6BD | ACP TE | Acyl-ACP thioesterase | final step in de novo synthesis - | " |
| III | Palmitic acid | AX-147240313 | 7.84E-05 | 2.89E-03 | B02 | 2497726 | 17.654 | arahy.LCII34 | ACP TE | Acyl-ACP thioesterase | final step in de novo synthesis - terminates chain growth | " |
| III | Arachidic acid | AX-147244543 | 4.62E-06 | 7.29E-03 | B03 | 43677278 | 578.389 | arahy.KT8BMX | ACP TE | Acyl-ACP thioesterase | final step in de novo synthesis - terminates chain growth | Feng et al., 2018; Feng et al., 2017; Jones et al., 1995 |
| III | Unsaturated fatty acid | AX-147247670 | 2.33E-09 | 3.22E-06 | B04 | 72562327 | 843.687 | arahy.QR9F0J | ACP TE | Acyl-ACP thioesterase | final step in de novo synthesis - terminates chain growth | Feng et al., 2018; Feng et al., 2017; Jones et al., 1995 |
| III | Palmitic acid | AX-177640893 | 4.80E-07 | 1.77E-03 | B10 | 127038914 | 937.449 | arahy.L4EP3N | ACP TE | Acyl-ACP thioesterase | final step in de novo synthesis - terminates chain growth | " |
| IV | Palmitic acid | AX-176797560 | 2.34E-05 | 1.77E-03 | A04 | 6029070 | 561.751 | arahy.WX9SXY | ACP I | acyl carrier protein 1 | carrier of the growing fatty-acyl chain | Chen et al., 2011; Chen 2012 |
| IV | Linoleic acid | AX-147221650 | 2.52E-06 | 6.95E-03 | A05 | 7266706 | 43.462 | arahy.SNFJ12 | ACBD3 | acyl-CoA-binding domain 3 | Binds medium- and long-chain acyl-CoAs | Chapman and Ohlrogge, 2012; Li-Beisson et al., 2013; Hurlock et al., 2014 |
| IV | Gadoleic acid | AX-176818595 | 3.68E-08 | 1.35E-04 | B06 | 17396747 | 333.08 | arahy.IB7YTW | ACBD4 | acyl-CoA-binding domain-containing protein 4-like [Glycine max] | Binds medium- and long-chain acyl-CoAs | " |
| IV | Stearic acid | AX-177644201 | 6.23E-07 | 1.37E-03 | B08 | 111389374 | 190.097 | arahy.FIRH26 | FABP | UPF0678 fatty acid-binding protein-like protein | extra- and intracellular membrane transport of fatty acids | Kader, 1996; Arondel et al., 1990 |
| IV | Unsaturated fatty acid | AX-177637654 | 8.18E-08 | 2.04E-04 | B08 | 111377562 | 201.909 | arahy.FIRH26 | FABP | UPF0678 fatty acid-binding protein-like protein | extra- and intracellular membrane transport of fatty acids | " |
| IV | Palmitic acid | AX-177640958 | 7.42E-14 | 3.15E-11 | B09 | 40167055 | 180.24 | arahy.6BI416 | ACBD4 | acyl-CoA-binding domain-containing protein 4-like [Glycine max] | Binds medium- and long-chain acyl-CoAs | " |
| V | Linoleic acid | AX-147208865 | 1.80E-07 | 4.97E-04 | A01 | 5940154 | 68.91 | arahy.0JDQ22 | FAD8 | fatty acid desaturase 8 | unsaturated fatty acid biosynthesis | " |
| V | Palmitic acid | AX-147212708 | 1.36E-06 | 4.07E-04 | A02 | 2658782 | 528.065 | arahy.XVN2S5 | ACP desaturase | Acyl-[acyl-carrier-protein] desaturase | unsaturated fatty acid biosynthesis | " |
| V | Arachidic acid | AX-176802895 | 4.16E-15 | 4.59E-11 | A04 | 98974824 | 832.816 | arahy.9ET73H | FAD8 | fatty acid desaturase 8 | unsaturated fatty acid biosynthesis | Phan et al., 2015; Chen et al., 2011; Gibson et al., 1994; Berberich et al., 1998 |
| V | Behenic acid | AX-147223115 | 1.80E-08 | 6.63E-05 | A05 | 89921606 | 742.981 | arahy.ZRK7H3 | FAD6 | fatty acid desaturase 6 | unsaturated fatty acid biosynthesis | " |
| V | Palmitic acid | AX-176823392 | 1.07E-04 | 3.45E-03 | A05 | 90972150 | 303.763 | arahy.ZRK7H3 | FAD6 | fatty acid desaturase 6 | unsaturated fatty acid biosynthesis | " |
| V | Linoleic acid | AX-147234396 | 4.89E-26 | 5.40E-22 | A09 | 114692575 | 516.677 | arahy.42CZAS | FAD2 | fatty acid desaturase 2 | unsaturated fatty acid biosynthesis | Dar et al., 2017; Bhunia et al., 2016; P Zhang et al., 2012 |
| V | Oleic acid | AX-147234396 | 1.18E-27 | 1.31E-23 | A09 | 114692575 | 516.677 | arahy.42CZAS | FAD2 | fatty acid desaturase 2 | unsaturated fatty acid biosynthesis | " |
| V | Palmitic acid | AX-147234396 | 4.54E-15 | 8.35E-12 | A09 | 114692575 | 516.677 | arahy.42CZAS | FAD2 | fatty acid desaturase 2 | unsaturated fatty acid biosynthesis | " |
| V | Unsaturated fatty acid | AX-176794805 | 3.08E-10 | 3.41E-06 | A10 | 28633929 | 462.929 | arahy.H654N0 | ACP desaturase | Acyl-[acyl-carrier-protein] desaturase | unsaturated fatty acid biosynthesis | Hernandez et al., 2019; Fox et al., 2004 |
| V | Palmitic acid | AX-177639949 | 2.80E-07 | 5.06E-05 | A10 | 2893055 | 322.423 | arahy.C923ZL | SLD1 | sphingolipid delta desaturase | unsaturated fatty acid biosynthesis | Chi et al., 2011; Ryan et al. 2007 |
| V | Palmitic acid | AX-147242934 | 1.80E-05 | 1.77E-03 | B03 | 1074146 | 657.517 | arahy.Y29W5P | STE | delta(7)-sterol-C5(6)-desaturase-like protein | unsaturated fatty acid biosynthesis |  |
| V | Palmitic acid | AX-147255424 | 1.26E-06 | 4.63E-03 | B07 | 27914176 | 443.178 | arahy.BC0JZ1 | FAD8 | fatty acid desaturase 8 | unsaturated fatty acid biosynthesis | " |
| V | Palmitic acid | AX-177639725 | 9.82E-05 | 3.32E-03 | B09 | 2036451 | 331.364 | arahy.GU7MJ6 | FAD5 | fatty acid desaturase 5 | unsaturated fatty acid biosynthesis | Bhunja et al., 2016; Phan et al., 2015; Zhang et al., 2012; Chen et al., 2011 |
| VI | Palmitic acid | AX-176803680 | 9.68E-05 | 3.29E-03 | A01 | 37436705 | 361.398 | arahy.MWEA9C | ACX1 | acyl-CoA oxidase 1 | metabolism [beta-oxidation] of unsaturated fatty acids | Cantu et al., 2012 |
| VI | Unsaturated fatty acid | AX-176794786 | 5.12E-28 | 5.66E-24 | A01 | 101033367 | 405.254 | arahy.YC35GQ | BCE2 | Lipoamide acyltransferase component of branched-chain alpha-keto acid dehydrogenase complex | beta-oxidation of straight-chain fatty acids to lipoic acid |  |
| VI | Linoleic acid | AX-147211846 | 6.01E-06 | 5.10E-03 | A01 | 96848733 | 118.266 | arahy.N8MRLM | CER1 | Fatty acid hydroxylase superfamily | hydroxylation of free fatty acids to form sphingolipids | " |
| VI | Palmitic acid | AX-147212208 | 5.77E-06 | 1.33E-03 | A02 | 453216 | 553.747 | arahy.GX90LB | CER1 | Fatty acid hydroxylase superfamily | hydroxylation of free fatty acids to form sphingolipids | " |
| VI | Palmitic acid | AX-147217615 | 1.06E-06 | 2.93E-03 | A03 | 118257901 | 793.297 | arahy.ZY648K | CER1 | fatty acid amide hydrolase-like [Glycine max] | hydroxylation of free fatty acids to form sphingolipids | " |
| VI | Unsaturated fatty acid | AX-176814481 | 4.01E-12 | 1.11E-08 | A03 | 119564950 | 508.316 | arahy.ZY648K | CER1 | fatty acid amide hydrolase-like [Glycine max] | hydroxylation of free fatty acids to form sphingolipids | Lee et al., 2015; Smith et al., 2003 |
| VI | Palmitic acid | AX-147219799 | 1.91E-05 | 1.77E-03 | A04 | 28582103 | 948.742 | arahy.BS3FTI | FAR3 | fatty acyl-CoA reductase 3-like [Glycine max] | reduction of fatty acyl-CoA to fatty alcohols | Doan et al., 2009; Rowland et al., 2006 |
| VI | Palmitic acid | AX-147219434 | 7.14E-06 | 1.49E-03 | A04 | 9842373 | 121.116 | arahy.407LUN | CER1 | fatty acid hydroxylase 1 | hydroxylation of free fatty acids to form sphingolipids | Lee et al., 2015; Smith et al., 2003 |
| VI | Oleic acid | AX-176815173 | 1.23E-06 | 2.72E-03 | A05 | 105438012 | 543.222 | arahy.531WVK | CER1 | Fatty acid hydroxylase superfamily | hydroxylation of free fatty acids to form sphingolipids | " |
| VI | Palmitic acid | AX-147226322 | 4.09E-05 | 1.98E-03 | A06 | 105426750 | 554.484 | arahy.531WVK | CER1 | Fatty acid hydroxylase superfamily | hydroxylation of free fatty acids to form sphingolipids | " |
| VI | Palmitic acid | AX-147233144 | 4.43E-11 | 1.25E-08 | A09 | 25073192 | 879.537 | arahy.RDH47T | ACX5 | acyl-CoA oxidase 5 | beta-oxidation of unsaturated fatty acids | Cantu et al., 2012 |
| VI | Linoleic acid | AX-176820362 | 1.99E-06 | 4.41E-03 | B03 | 2124257 | 222.343 | arahy.NJ7QKI | FAR | fatty acyl-CoA reductase | reduction of fatty acyl-CoA to fatty alcohols | " |
| VI | Palmitic acid | AX-147243024 | 8.01E-05 | 2.94E-03 | B03 | 2014046 | 332.554 | arahy.NJ7QKI | FAR | fatty acyl-CoA reductase | reduction of fatty acyl-CoA to fatty alcohols | " |
| VI | Palmitic acid | AX-147247065 | 4.08E-05 | 1.98E-03 | B04 | 10497585 | 828.215 | arahy.8UG7AL | CER1 | fatty acid hydroxylase 1 | hydroxylation of free fatty acids to form sphingolipids | " |
| VI | Palmitic acid | AX-177641788 | 1.81E-07 | 9.50E-05 | B08 | 5477774 | 584.395 | arahy.VAJ8HR | FAR5 | probable fatty acyl-CoA reductase 5-like [Glycine max] | reduction of fatty acyl-CoA to fatty alcohols | Doan et al., 2009; Rowland et al., 2006 |
| VI | Palmitic acid | AX-176820256 | 8.17E-05 | 2.98E-03 | B09 | 132164709 | 90.351 | arahy.RNBV6A | CFA | Cyclopropane-fatty-acyl-phospholipid synthase | adds mythelene to unsaturated fatty acids | Bao et al., 2002 |
| VI | Palmitic acid | AX-147261782 | 7.52E-05 | 2.84E-03 | B09 | 131670685 | 0 | arahy.08UV73 | CFA | Cyclopropane-fatty-acyl-phospholipid synthase | adds mythelene to unsaturated fatty acids | Bao et al., 2002 |

These results suggest the involvement of more alleles in determining fatty acid composition than previously reported in peanut. For years, peanut scientists have believed the oleic trait to be controlled by mutations in two homeologous *Arachis hypogaea* FAD2 genes (ahFAD2) (Chu et al. 2009; Jung et al. June 2000; Lopez et al. 2002; Pandey et al. 2014; Patel et al. 2004; Wang et al. 2011). Thus, the genetic control of this trait is more complex than previously thought.

### Associations with variants proximal to 55 unique genes encoding ACCase subunits and different FAS II enzyme components

Based on the gene(s) within 1000 kbp of the detected associations, we grouped the associations into six groups following the roles the genes have been reported to play in fatty acid biosynthesis and metabolism. **Group I** neighboring genes encode acetyl-CoA carboxylase carboxyl transferase subunits alpha and beta (**Table 3**). The SNPs in this group are associated with saturated and unsaturated fatty acid content (**Figure 2: D-E**). The SNP associated with saturated fatty acid content is only ∼7 kbp from gene arahy.15M9SP, which encodes Acetyl-CoA carboxylase carboxyl transferase subunit beta (accB). Acetyl-CoA carboxylase catalyzes the first committed step in de novo fatty acid synthesis (Freiberg et al. 2004).

**Figure 1.**
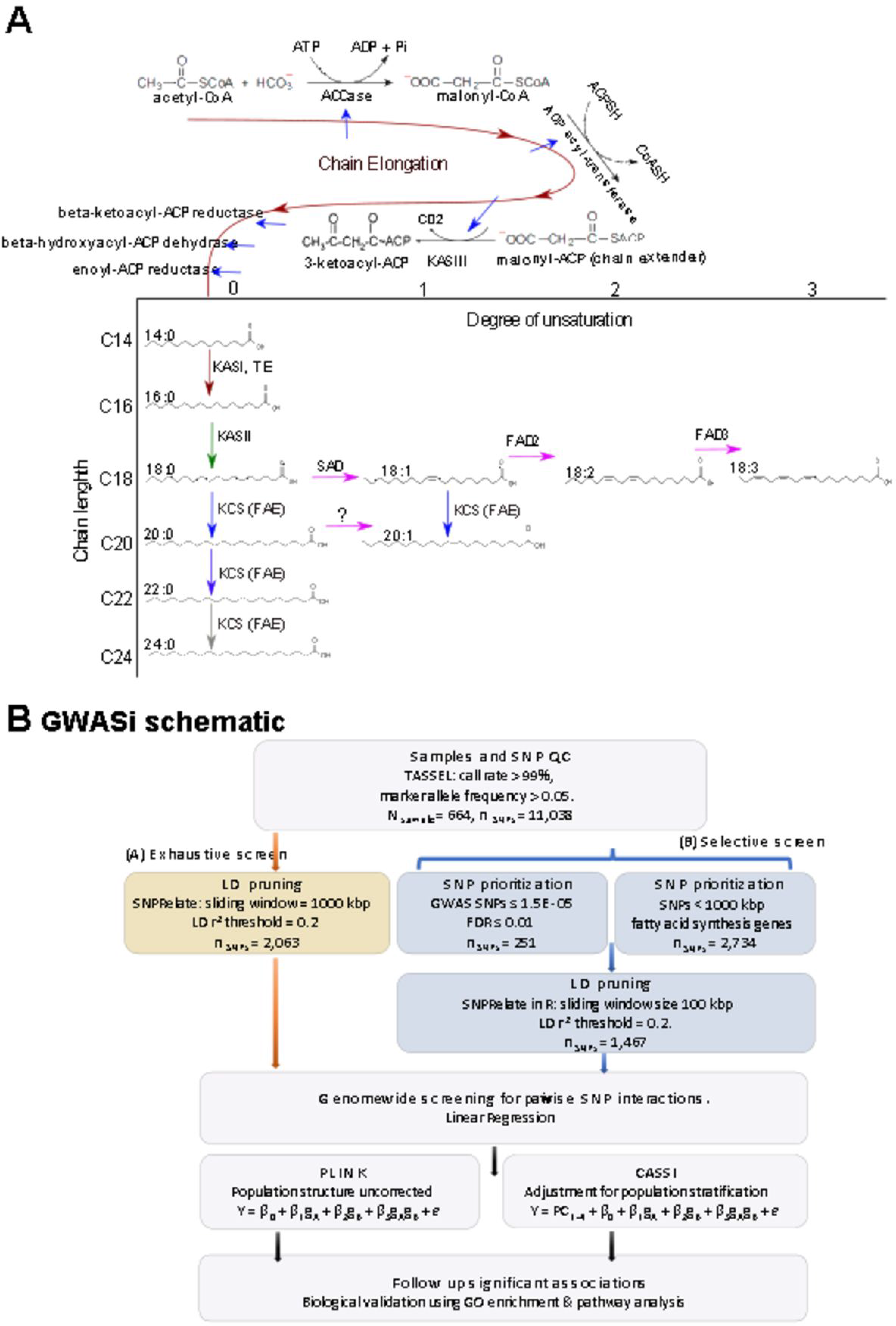
A. De novo fatty acid synthesis and post-synthesis modification in *A. hypogaea*. **B.** Schematic for Genome-wide association interactions screening approaches employed in the study. Inferred fatty acid synthesis pathway was modified from Carlson et al. (2019).

**Figure 2.**
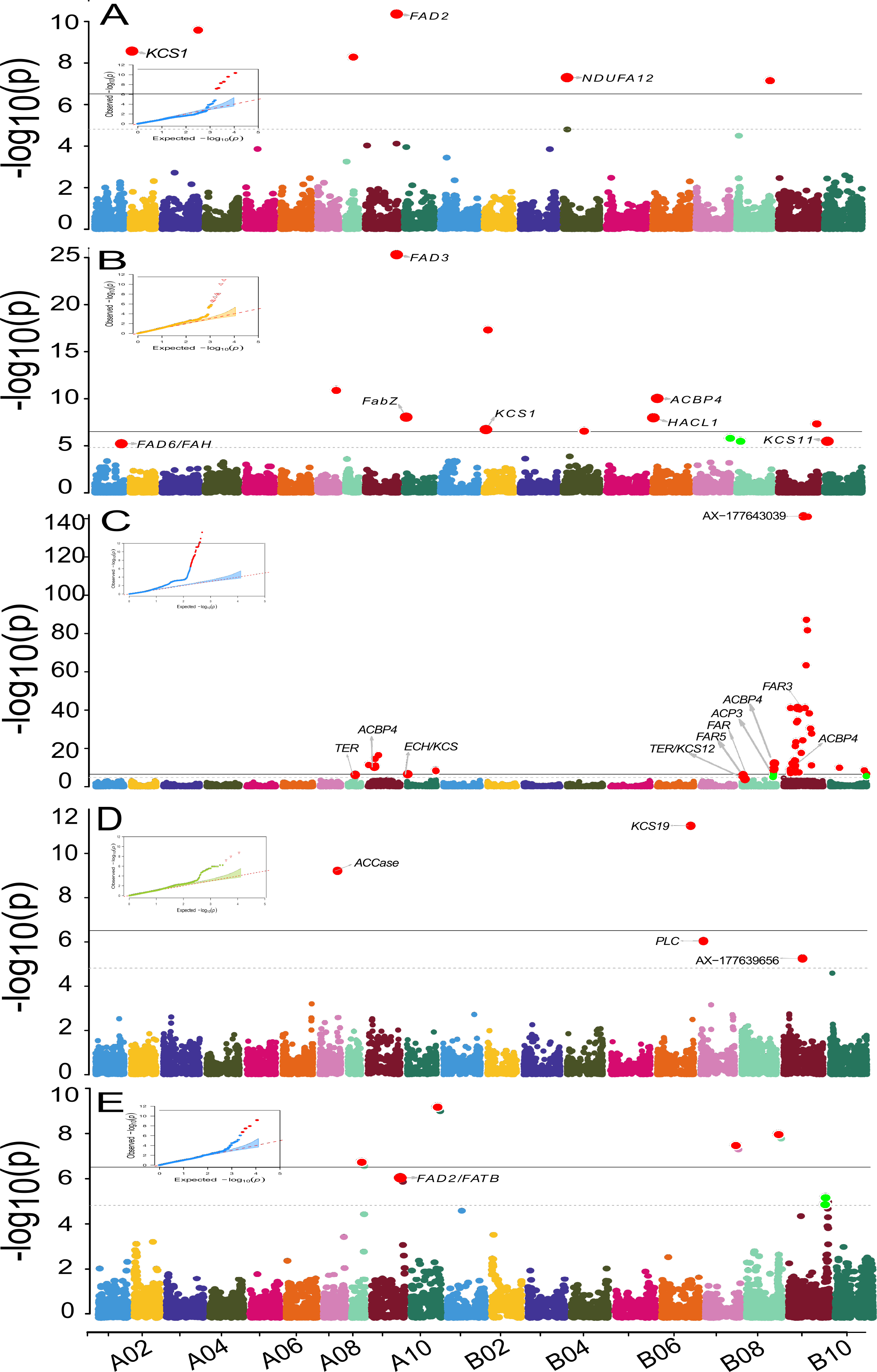
Manhattan plots of GWAS results for: **A.** Oleic acid content, **B.** Linoleic acid content, **C.** Palmitic acid content, **D.** Saturated fatty acid content, and **E.** Unsaturated fatty acid content. Genes with known fatty acid biosynthesis function are annotated atop lead SNPs if they occur within 1000 kbp of the SNP. QQ plots are enclosed.

Associations in **Group II** neighboring genes encode enzymes catalyzing condensation reactions for chain growth via the addition of two carbon atoms to the growing fatty-acyl chain, from malonyl-CoA to a 16-C fatty-acyl compound. We identified 16 unique peanut genes encoding 3- oxoacyl-[acyl-carrier-protein] synthase 3 (KAS III), 3-ketoacyl-CoA synthase 1 (KCS1), KCS2, KCS4, KCS11, KCS12, KCS19, chalcone synthase (CHS), 3-hydroxyacyl-[acyl-carrier-protein] dehydratase (FabZ), and 3-hydroxyacyl-CoA dehydrogenase family protein (HADH) (**Table 3**). KAS III catalyzes the first condensation reaction in FAS II elongation cycle (Choi et al. 2000; White et al. 2005). KCS1 is a long-chain fatty acid elongase enzyme belonging to the Ketoacyl synthase group of enzymes – particularly Ketoacyl synthase 2 enzyme family. Ketoacyl synthase enzymes (KSs) catalyze condensing reactions that produce 3-ketoacyl-CoA by combining malonyl-CoA with acyl-CoA. KSs will also produce 3-ketoacyl-ACP when an acyl-acyl carrier protein (acyl-ACP) is joined to malonyl-CoA (Blacklock and Jaworski 2006; Chen 2012; Chen et al. 2011). This reaction is a key step in the fatty acid synthesis cycle, contributing to de novo chain elongation by adding two carbon atoms to growing acyl chains. Five different Ketoacyl synthase families have been characterized with respect to their characteristic primary structures – KS1, KS2, KS3, KS4, and KS5 (Chen 2012; Chen et al. 2011). KS1 family enzymes are predominantly 3-ketoacyl-ACP synthase III and its variants produced almost exclusively by bacteria with a few eukaryota exceptions (Chen et al. 2011). KS2 enzymes are long-chain fatty acid elongases produced by plants while KS3, KS4, and KS5 are mainly large bacterial and eukaryotic families. The KS3 family is predominantly 3-ketoacyl-ACP synthase I and II while the majority of the KS4 family is composed of chalcone synthases (Chen et al. 2011). With the exception of arachidic and lignoceric acid content, the remaining saturated fatty acid components are associated with at least one SNP within 1000 kbp of six KCS genes (**Figure 2**, **Figure 3**, **Table S2**). The SNP AX-147247147, located ∼ 220 kbp from a Chalcone synthase gene, *arahy.YUL6YW,* is associated jointly with linoleic and oleic acid content (**Figure 2: A-B**). This SNP had the overall effect of lowering linoleic and increasing the oleic acid content of all genotypes carrying the favorable allele (**Figure 4: A-B**, **Figure 5: E**). *FabZ* is one of two forms in which β- hydroxyacyl-(acyl carrier protein, ACP) dehydratase exists along with *FabA*. Both isoforms catalyze the third step in the fatty acyl elongation cycle involving the dehydration of (3R)- hydroxyacyl-ACP to form trans-2-acyl-ACP. *FabA* also catalyzes the isomerization of trans-2- to cis-3-decenoyl-ACP, an essential step in unsaturated fatty acid biosynthesis (Heath and Rock 1996; Shimakata and Stumpf 1982). 3-hydroxyacyl-CoA dehydrogenase (HADH) is an oxidoreductase enzyme that catalyzes the third step in fatty acid synthesis. During this beta oxidation step, the hydroxyl in the growing fatty acyl chain is converted to 3-ketoacyl CoA (Cantu et al. 2012).

**Figure 3.**
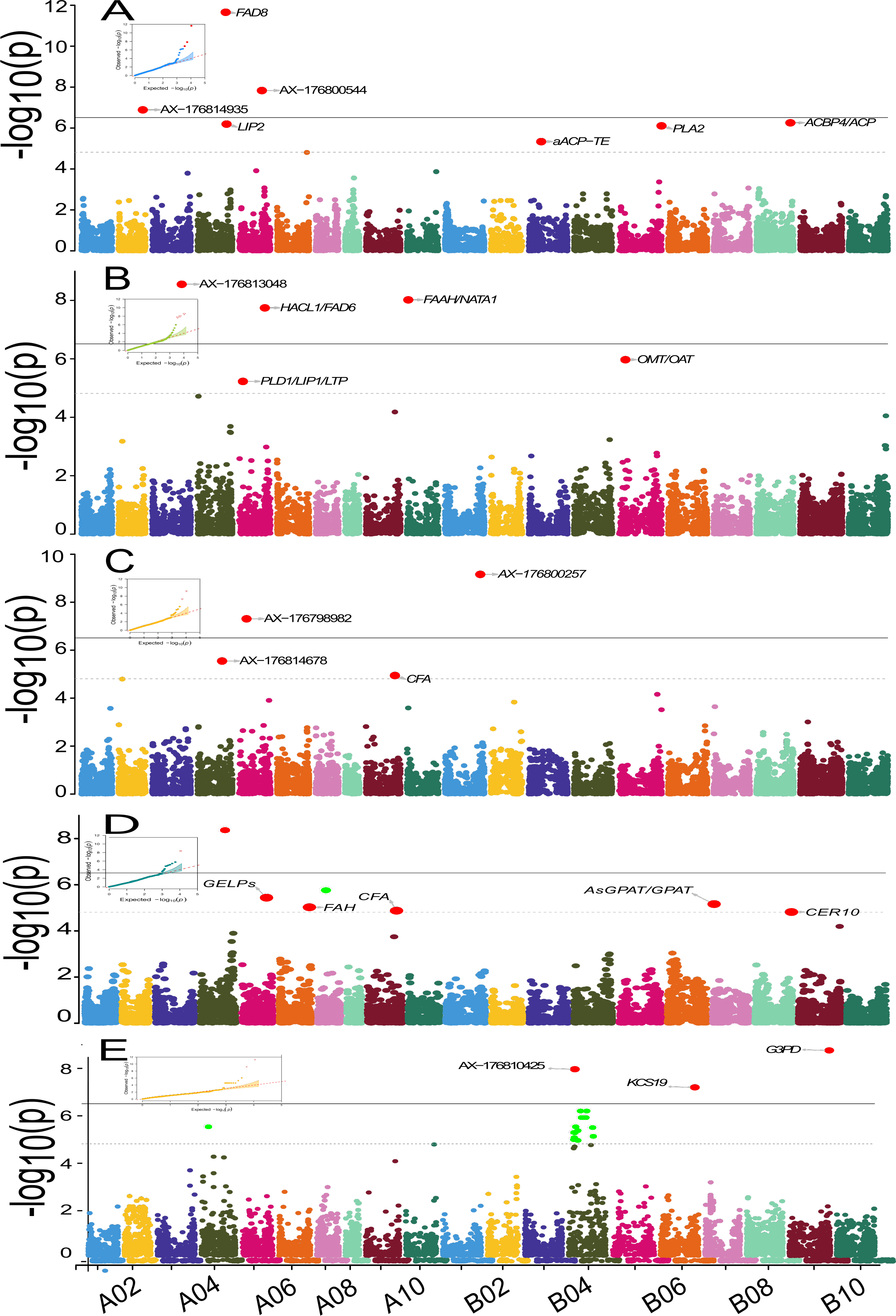
Manhattan plots of GWAS results for: A. Arachidic acid content, **B.** Behenic acid content, **C.** Lignoceric acid content, **D.** Gadoleic acid content, and **E.** Stearic acid content. Genes with known fatty acid biosynthesis function are annotated atop lead SNPs if they occur within 1000 kbp of the SNP. QQ plots are enclosed.

**Figure 4.**
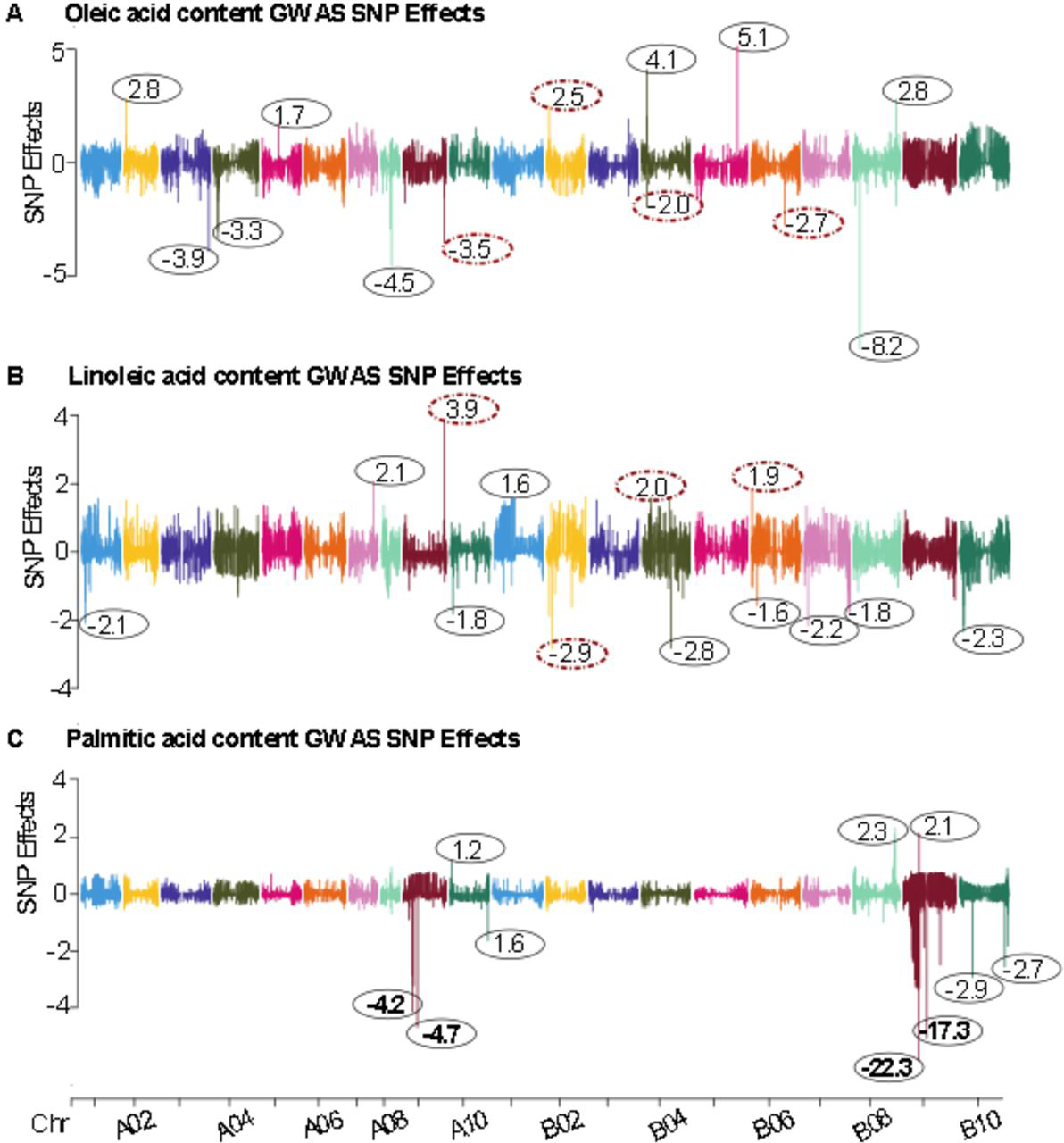
SNP effects for GWAS on: **A.** Oleic acid content, **B.** Linoleic acid content, and **C.** Palmitic acid content. SNPs with joint effects on Oleic and Linoleic acid content are shown with their effect sizes in red-colored oval rings with broken lines. SNPs with very large effect sizes on Palmitic acid content are shown with effect sizes in bold.

Associations in **Group III** neighboring genes encode Acyl-ACP thioesterases (**Table 3**). The SNPs in this group are associated with palmitic, saturated, unsaturated fatty acid content (**Figure 2: C-E**), and arachidic acid (**Figure 3. A**). The SNP AX-176819119 (B02) is associated with palmitic acid content. This SNP occurs within an Acyl-ACP thioesterase encoding peanut gene *arahy.ZKU6BD* (**Table 3**), making it a strong candidate for functional validations. Acyl-ACP thioesterase (TE) is a key FASII enzyme that determines the carbon chain length of fatty acyl compounds by catalyzing the hydrolysis of thioester bonds to terminate de novo fatty acid synthesis (Feng et al. 2017; Feng et al. 2018; Jones et al. 1995). Most TEs play a key part in recognizing palmitoyl (16:0) and oleoyl (18:1) substrates, and only a few specific TEs are responsible for the termination of acyl-ACP elongation at C8-C14 (Feng et al. 2017). In **Group IV**, we categorized associations of neighboring genes that encode acyl carrier protein 1 (ACP1), acyl-CoA-binding domain 3 (ACBD3), ACBD4, and fatty acid binding protein (FABP). These SNPs are associated with linoleic, gadoleic, unsaturated, palmitic, and stearic acid content (**Table 3**). These proteins bind medium- to long-chain acyl-CoAs and transport the growing fatty-acyl chain across membranes (Arondel et al. 1990; Chapman and Ohlrogge 2012; Chen 2012; Hurlock et al. 2014; Li-Beisson et al. 2013).

Following de novo synthesis, endoplasmic reticulum (ER) and chloroplast bound fatty acid desaturase (FAD) enzymes modify fatty-acyl compounds via the introduction of double bonds into the chain. We categorized associations proximal to fatty acid desaturase genes in **Group V** (**Table 3**). These SNPs are associated with oleic, linoleic, palmitic, unsaturated, arachidic, and behenic fatty acid content (**Figure 2: A-C, E and Figure 3: A-B**). Stearoyl-acyl carrier protein desaturase (SAD) catalyzes the desaturation of stearic acid (C18:0) to oleic acid (C18:1), while both FAD2 and FAD6 catalyze the desaturation of oleic acid to linoleic acid (18:2) in the ER and plastids, respectively. FAD3 and FAD7/FAD8 catalyze the desaturation of linoleic acid to γ-linolenic acid (C18:3, n6) in the ER and plastids, respectively (Bhunia et al. 2016; Zhang et al. 2012). ER bound FAD2 and FAD3 genes also act on extra chloroplast lipids (Los and Murata 1998; Shanklin and Cahoon 1998). Oleic, linoleic, and palmitic acid content jointly associated with the SNP, AX-147234396 (A09; P values = 1.18E-27, 4.89E-26, 4.54E-15). This SNP is ∼ 516 kbp from a fatty acid desaturase (FAD) 2 gene, arahy.42CZAS. We detected associations within 1000 kbp of FAD2, FAD5, FAD6, FAD8, Acyl-[acyl-carrier-protein] desaturase, sphingolipid delta desaturase (SLD1), and delta(7)-sterol-C5(6)-desaturase-like protein (STE) (**Table 3**).

In the final group, **Group VI**, we categorized associations within 1000 kbp of genes involved in fatty acid metabolism. These SNPs are associated with linoleic, palmitic, and unsaturated fatty acid content (**Table 3**, **Figure 2: B,C,E**). We evaluated eight peanut genes in this group which encoded for acyl-CoA oxidase 1 (ACX1), ACX5, Cyclopropane-fatty-acyl-phospholipid synthase (CFA), fatty acyl-CoA reductase (FAR3), FAR3, FAR5, Lipoamide acyltransferase component of branched-chain alpha-keto acid dehydrogenase complex (BCE2), fatty acid hydroxylase superfamily (CER), and CER1. These genes are involved in the beta-oxidation of unsaturated fatty acids, addition of a methylene bridge to unsaturated fatty acids, reduction of fatty acyl-CoA to fatty alcohols, beta-oxidation of straight-chain fatty acids to lipoic acid, and in the hydroxylation of free fatty acids to form sphingolipids (Bao et al. 2002; Cantu et al. 2012; Doan et al. 2009; Lee et al. 2015; Rowland et al. 2006; Smith et al. 2003).

### A delicate balance: Breeding for a high Oleic to Linoleic acid (O/L) ratio

Breeding for long shelf-life, high nutritional quality, and flavor requires a delicate balance of fatty acid composition in the oil. Unlike oleic acid, the human body is incapable of producing linoleic acid to meet its functional needs. This makes linoleic acid an essential nutrient, supplied solely through the diet or as supplements. However, high amounts of linoleic acid in peanut seed oil increases susceptibility to oxidative rancidity which reduces the shelf-life and over time can offset flavor. Conversely, high amounts of oleic acid content extends shelf-life due to its relatively high oxidative stability compared to linoleic acid (Pandey et al. 2014). Thus, peanuts and oils with high oleic to linoleic acid ratio are desirable and fetch a premium market price. Additionally, the consumption of oils rich in monounsaturated fatty acids such as high oleic, has been linked with overall better cardiovascular health, tumor suppression, and protection from inflammatory diseases (O’Byrne et al. 1997; YAMAKI et al. 2005).

We detected a strong negative correlation (r^2^ = 0.91) between oleic and linoleic acid content in seed oils (**Figure S1. A**). Strikingly, oleic acid was more abundant than linoleic acid in all phenotyped lines except for PI 386350 (**Table 1**). The ratio of oleic to linoleic acid content (O/L) ranged from 2.24 to 28.59 in commercial cultivars and from 0.94 to 6.37 in PI accessions except for PI 461451 which had an O/L ratio of 28.79 comparable to the best performing cultivars. Among cultivars, Red River Runner / PI 665474 (Melouk et al. 2013), Florida-07 / PI 652938 (Gorbet and Tillman 2009), Florida Fancy / PI 654368 (Branch 2007), FloRun 107 / PI 663993 (Tillman and Gorbet 2015), and NM309-2 / PI 670460 (Puppala and Tallury 2014), had the highest O/L ratio, ranging from 7.14 to 28.6. While on the low end, O/L ranged from 2.39 to 3.13 for Georgia-06G / PI 644220 (Branch 2007), Bailey / PI 659502 (Isleib et al. 2011), and Jupiter (Anon. 2000).

High O/L cultivars are specifically bred and selected to carry mutations suppressing the expression of two homeologous *Arachis hypogaea* FAD2 genes (here on abbreviated as *ahFAD2A* on chromosome A09 and *ahFAD2B* on B09) (Lopez et al. 2002; Pandey et al. 2014; Patel et al. 2004; Wang et al. 2011). (Chu et al. 2009) showed that two alleles of *ahFAD2B* are responsible for the high O/L trait in U.S. grown peanuts.

We detected 10 SNPs jointly associated with oleic and linoleic acid content at different positions in the genome (**Table 2**). For all 10 SNPs, the favorable allele had opposite effects on the oleic acid and linoleic acid content of genotypes with these alleles (**Figure 4: A-B**, **Figure 5: A-J**). Six of the 10 SNPs reduced the overall oleic content while increasing linoleic acid content for genotypes with the favorable allele. Only four SNPs produced the desired effect of increasing oleic content while decreasing linoleic acid content (**Figure 4: A-B**, **Table 2**).

The SNP AX-147234396 is jointly associated with oleic, linoleic, and palmitic acid content (**Figure 4: A-C**, **Figure 5. A**). Genotypes with the favorable allele at this position had a small marginal increase in palmitic acid content (0.71), but with a marked decrease in oleic acid content (-4.51) and a high increase in linoleic acid content (3.89) (**Figure 4: A-C**, **Table 2**). Although the causal locus underlying this behavior is not fully validated, this SNP is ∼ 516 kbp from the *ahFAD2A* gene, *arahy.42CZAS* on chromosome A09. It is thus likely this gene, ah*FAD2A,* affects fatty acid composition in more ways than previously reported, functioning to control the concentration of palmitic acid content as well as oleic and linoleic acid content in *A. hypogaea*. (Chu et al. 2007)reported the 448G<A base substitution in *ahFAD2A* is prevalent in *A. hypogaea* subsp. *hypogaea* accessions in the U.S. mini core collection. This observation is likely more common and applies more widely with subsp. *hypogaea* accessions across the entire U.S core collection. It is noteworthy that most of the market share of peanuts grown in the USA are of subspecies *hypogaea* (runner market class) in spite of there being less accessions classified in this subspecies in the entire peanut germplasm collection.

Unlike *ahFAD2A*, the mutation in *ahFAD2B* is quite rare and this could explain the lack of a strong association signal in this region for oleic and linoleic acid, in spite of the well-characterized causal mutation in this region. Nevertheless, the region shows a strong association with palmitic acid content (**Figure 2. C**, **Figure 4. C**). A known weakness of GWAS is its lack of power to detect rare alleles that are involved in natural variation thus contributing to missing heritability (Brachi et al. 2011; Young 2019; Zuk et al. 2012).

Since saturated fatty acids are associated with overall poor cardiovascular health, we highlight a SNP detected with joint effects on the total saturated and unsaturated fatty acid content (**Figure 5. K**, **Table 2**). This SNP, AX-176791407, is located on chromosome A09 ∼ 851 kbp from an Acyl-ACP thioesterase (TEs) encoding gene, arahy.75YJ3Z. Genotypes carrying the desirable allele had a unit increase in the amounts of unsaturated and a unit decrease in the overall saturated fatty acid content (**Table 2**).

The overall abundance of saturated fatty acids in peanut seed oil ranged from 9.8 to 109.3, with a median value of 21.3. Palmitic acid was the most abundant saturated fatty acid. We detected several SNPs, both in LD and not, on chromosomes A09 and B09 with effect sizes ranging -4.2 to -22.3. Thus, in addition to being key targets for breeding high O/L peanuts, chromosomes A09 and B09 can serve as key targets for selecting overall high amounts of unsaturated fatty acids compared to saturated fatty acids (**Figure 4: A-C**).

### GWASi: contribution of SNP-SNP interaction (epistasis) to variation in fatty acid composition

Epistasis is when two or more different genes contribute to a single phenotype and their effects are not merely additive (Fisher 1918). Gene interactions can thus create new or modify existing phenotypes. In canonical association studies, such interactions are usually missed although a significant portion of the genetic variability of a trait can be a result of gene interactions.

We implemented an exhaustive and a selective screening approach to investigate the contribution of epistatic interactions on the variation of fatty acid composition (**Figure S1. B**). We detected relatively fewer interactions using CASSI (13,991 and 7,682 SNP pairs) compared to PLINK (40,378 and 24,168 SNP pairs) using the exhaustive and selective approaches, respectively.

CASSI allows for fitting PCA to correct for stratification while PLINK fits a regression model without covariates and thus does not account for confounding effects which might explain the inflated results between the two analyses. Here we present and discuss results from the selective strategy implemented in CASSI (**Table 4**).

**Table 4.** Summary of GWASi interactions detected via a selective epistasis scan in CASSI.

| Trait | GWAS pairs | GWAS x non-GWAS | Non-GWAS pairs | Total Interacting pairs |
| --- | --- | --- | --- | --- |
| Oleic acid content | 0 | 4 | 1,007 | 1,011 |
| Linoleic acid content | 0 | 9 | 1,238 | 1,247 |
| Gadoleic acid content | 0 | 0 | 0 | 0 |
| Palmitic acid content | 18 | 309 | 2,699 | 3,026 |
| Stearic acid content | 1 | 33 | 220 | 254 |
| Arachidic acid content | 0 | 0 | 77 | 77 |
| Behenic acid content | 0 | 2 | 3 | 5 |
| Lignoceric acid content | 0 | 0 | 0 | 0 |
| Saturated fatty acid content | 4 | 89 | 821 | 914 |
| Unsaturated fatty acid content | 6 | 112 | 1,030 | 1,148 |

With the exception of gadoleic and lignoceric acid content, all traits had significant SNP interactions exceeding a stringent Bonferroni threshold p-value < 4.65E-08 (**α** = 5%). Some SNPs had multiple interactions; thus, we selected the most significant SNP pair (lowest P value) as a representative of the interacting region to visualize genome-wide interactions in a circos plot (**Figure 6**).

**Figure 5.**
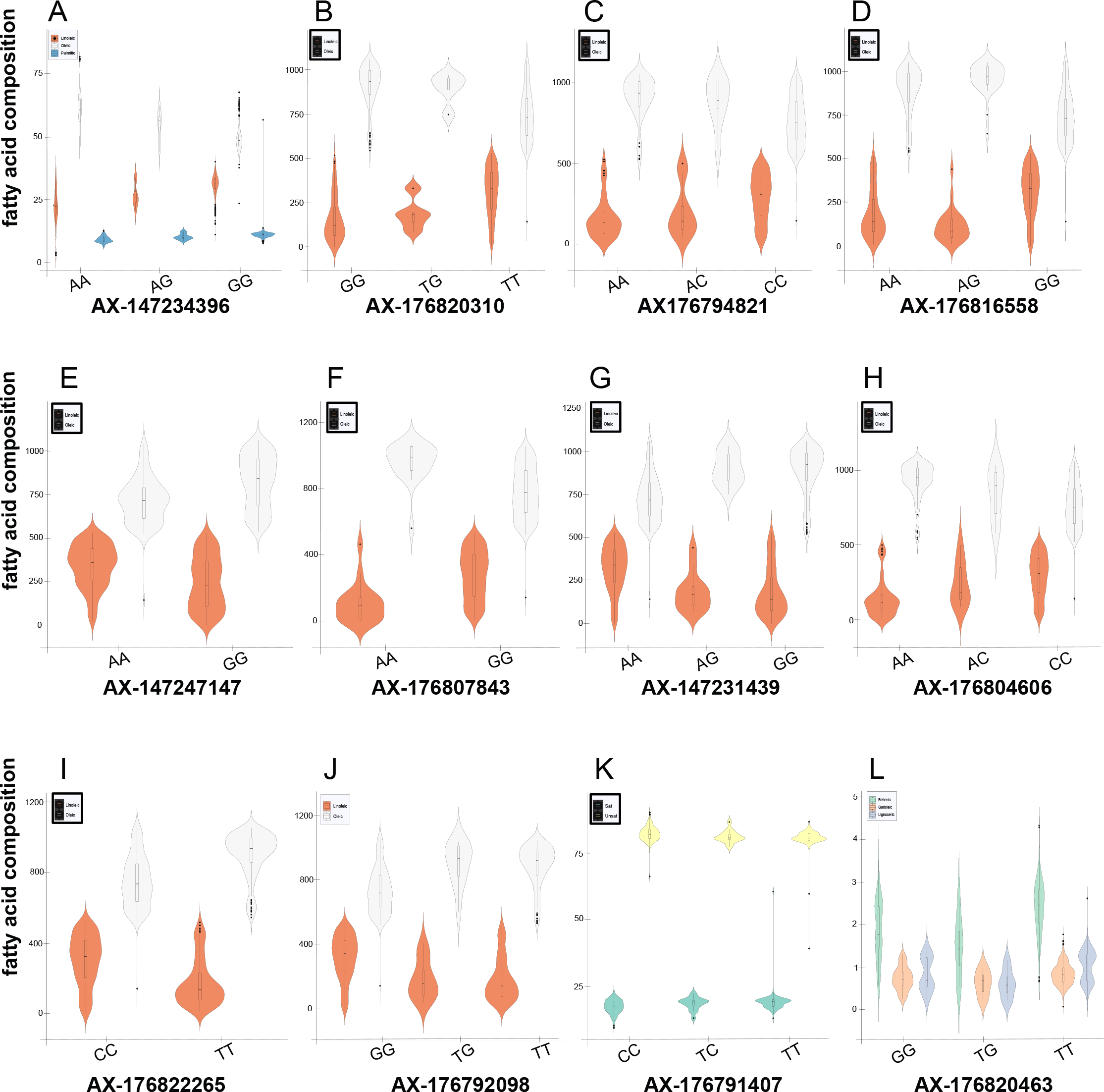
SNPs with apparent pleiotropic effects on fatty acid composition. The violin plots show the allele combinations in the population and their effect sizes on fatty acid abundance. **A.** SNP with joint effects on Oleic, Linoleic, and Palmitic acid content. **B-J.** SNPs with joint effects on Oleic and Linoleic acid content. **K.** SNP with joint effects on Saturated and Unsaturated fatty acid content. **L.** SNP with joint effects on Behenic, Gadoleic, and Lignoceric acid content.

**Figure 6.**
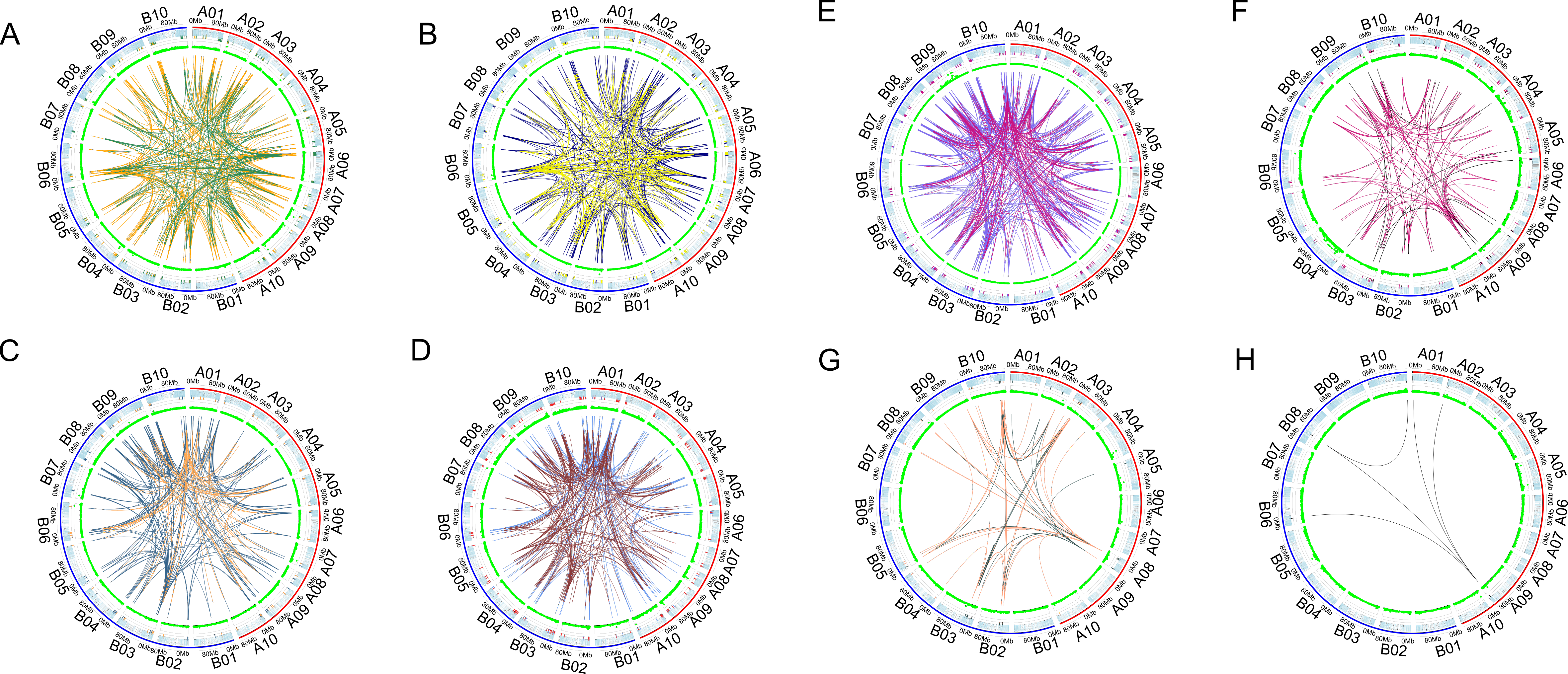
Circos plots showing epistatic interactions detected across the genome for different fatty acid components: **A.** Oleic acid content – the orange and green links show interactions with positive and negative effects, respectively; **B.** Linoleic acid content – the navy-blue and yellow links represent positive and negative effect interactions; **C.** Saturated fatty acids – green and coral links represent positive and negative interactions; **D.** Unsaturated fatty acids – brown and light-blue links represent positive and negative interactions; **E.** Palmitic acid content – purple and red-violet links represent positive and negative effect interactions; **F.** Stearic acid content – pink and black links represent positive and negative effect interactions; **G.** Arachidic acid content – salmon and dark-cyan links represent positive and negative interactions; **H.** Behenic acid content – black links represent interacting regions across the genome with negative effects on trait abundance. In the circos plot, the outer to innermost rings represent: cytobands for the **A (red)** and **B (blue) sub-genomes**; genome-wide SNPs with horizontal colored bars representing interacting SNP pairs; **GWAS results (light green)**; and **links connect interacting SNPs/regions**.

For major traits – oleic, linoleic, and palmitic acid, as well as trait totals –saturated and unsaturated fatty acid content, each of the 20 chromosomes had significant interactions with other chromosomal regions (**Figure 6: A-E**). Many of the interacting regions were previously undetected in GWAS (**Table 4**). No significant interactions were detected on chromosome B07 for stearic acid, while for arachidic acid, interactions were absent for A04, A06, A07, A09, B01, B06, and B07 (**Figure 6: F-G**). All five interactions contributing to variation in behenic acid content had negative effects on trait concentration. A SNP on chromosome A09 interacted with SNPs on chromosomes A01, A02, B06, and B08 to explain the variability in behenic acid content (**Figure 6. H**). Overall, palmitic acid content had the most interacting SNP pairs (3,026) contributing to trait variability followed by linoleic and oleic acid content with ∼ 50 % less interactions compared to palmitic acid at 1,247 and 1,011, respectively (**Table 4**).

These results, buoyed by the recent SNP genotyping of the core collection (Otyama et al. 2020), present new evidence for widespread interactions among SNPs with significant effects on the variation of fatty acid composition in this tetraploid species. This further supports the notion that more loci than previously detected play a significant role in determining fatty acid composition in peanut, with significant interactions among them. Previous studies have suggested epistasis as a contributor to fatty acid variation in peanut, particularly pointing to the significant interactions between the ahFAD2 genes (Barkley et al. 2013; Isleib et al. 2006).

### Enrichment analyses: Towards systems-level genetics controlling variation in seed fatty acid composition

Epistasis implies dependencies among genes in a network to maintain the phenotypic stability of critical cell components (Waddington 1942). This makes the identification of epistasis a key step to systems-level genetics. To understand the complexity of the biology underlying fatty acid composition in oil crops, we examined ontologies, gene families, pathways, and networks enriched in GWAS and GWASi gene lists.

We selected 627 SNP pairs from GWASi for which at least one of the significantly interacting SNP pairs is within 1000 kbp of a gene encoding proteins with known fatty acid biosynthesis function (**Table S4**). For these pairs, we identified 168 unique genes. To this list, we added 55 genes evaluated for their proximity to 100 SNPs identified via GWAS. Upon filtering for redundancies, we retained a total 177 unique genes for enrichment analysis. Enrichment was performed using LegumeMine, PeanutMine, and BLUEGENES hosted on the Legume Information System database (Berendzen et al. 2021; Dash et al. 2016; Smith et al. 2012).

Seven biological process GO terms, along with 39 gene families, were enriched in our analyses (**Figure S2**). Highly enriched gene families (P value < 4.0E-10) include: fatty acyl-CoA reductase 3-like, fatty acid hydroxylase superfamily, 3-ketoacyl-CoA synthase 11, 3-ketoacyl-CoA synthase 4, long chain alcohol oxidase FA02-like protein, acyl-ACP thioesterase, and acyl-[acyl-carrier-protein] desaturase (**Table S5**).

Additionally, we imported the list of genes identified via GWASi (168 unique genes) into STRING to confirm and validate known protein-to-protein interactions (Szklarczyk et al. 2016). First, we predicted gene families and proteins for these genes in *Glycine max*. From 168 unique *Arachis hypogaea* genes (hereafter referenced as “*arahy* genes”), we predicted 198 unique *Glycine max* protein identifiers (hereafter referenced as “*glyma genes*”) (**Table S5**). The resulting 198 *glyma* genes were imported into STRING for protein-to-protein interaction (PPI) analysis with settings: 1^st^ shell = none and 2^nd^ shell = none, to achieve full statistical validity. All p-values were corrected for multiple testing within each category using the Benjamin–Hochberg method.

Out of 198 gene IDs, 60 nodes (proteins) were returned in STRING. The predicted PPI network was significantly enriched for interactions (p-value < 1.0E-16) suggesting a level of biological connectedness among these proteins compared to all other proteins in the genome (**Figure 7**). Similarly, GO terms were significantly enriched for unsaturated fatty acid biosynthetic process (GO:0006636, p-value = 4.5E-05), fatty acid biosynthetic process (GO:0006633, p-value = 1.78E- 05), and oxidation-reduction process (GO:0055114, p-value = 0.00022). Pathway enrichment revealed 15 significantly enriched KEGG pathways from the network including: biosynthesis of unsaturated fatty acids (gmx01040, p-value = 8.12E-11), fatty acid elongation (gmx00062, p- value = 1.87E-10), fatty acid metabolism (gmx01212, p-value = 3.02E-21), and fatty acid biosynthesis (gmx00061, p-value = 3.22E-13) pathways (**Table S5**).

**Figure 7.**
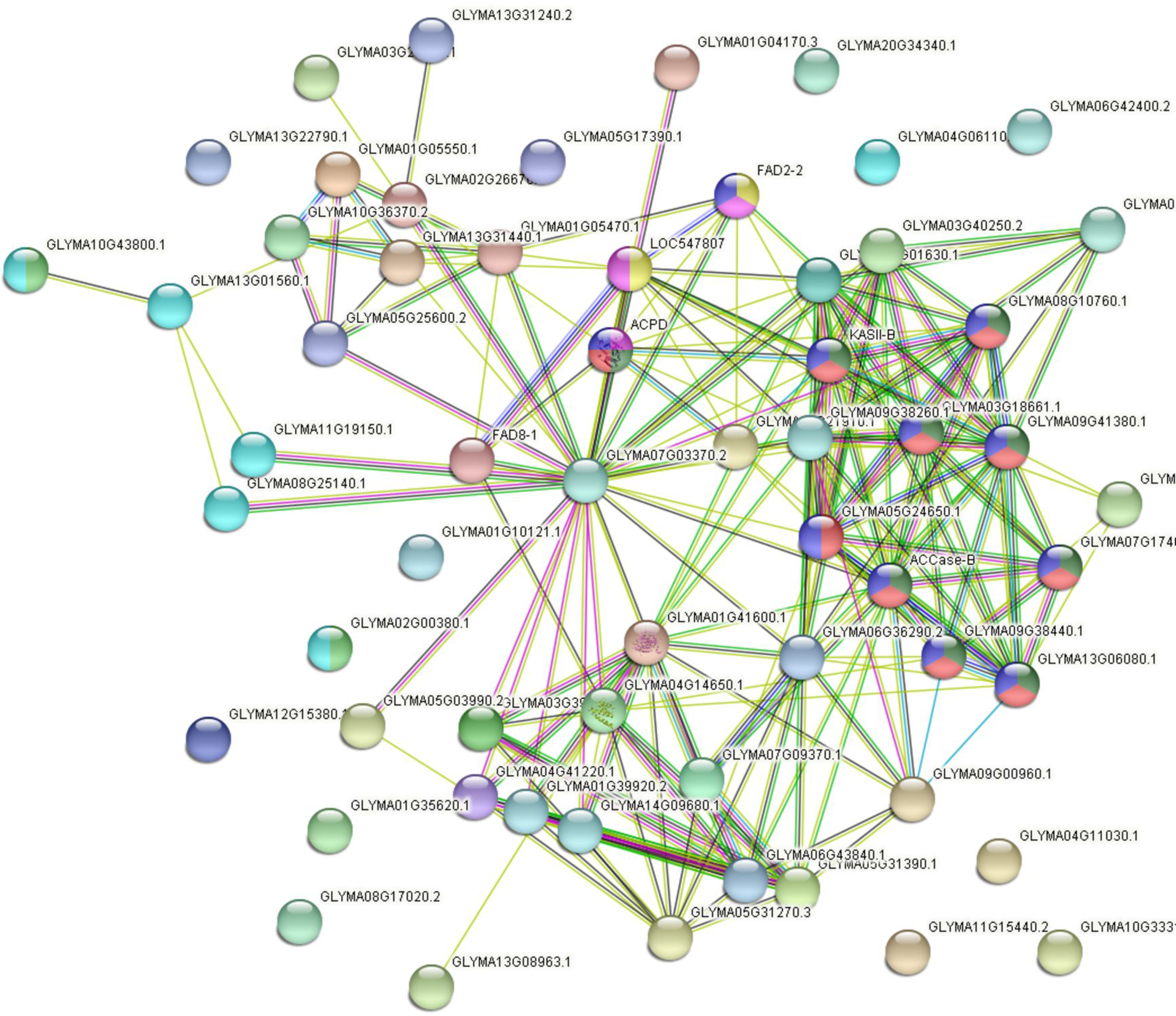
Protein-protein interactions of *Glycine max* genes predicted from *A. hypogaea* genes from GWASi. Nodes represent different genes, and the links represent interactions deduced from experimental evidence, text mining on publications, and or computationally derived evidence. Nodes are colored according to whether the represented genes are involved in fatty acid biosynthesis **(green),** lipid biosynthesis **(blue),** and/or fatty acid metabolism **(red).**

PPI evidence suggests a functional link between two Omega-6 fatty acid desaturases (FAD2) involved in the biosynthesis of 16:3 and 18:3 fatty acids (**Figure 7**). FAD2-2 (Glyma.03g144500) is thought to use cytochrome b5 as an electron donor in the endoplasmic reticulum whereas LOC547807 (Glyma.02g203300) uses ferredoxin as an electron donor to act on fatty acids esterified to galactolipids, sulfolipids and phosphatidylglycerol in the chloroplast (Heppard et al. 1996; Liu et al. 2015; Salimonti et al. 2020). Two acyl carrier proteins (Glyma.09g247100 and Glyma.09G014000) were also linked to these FAD2 genes (**Figure 7**) suggesting an intricate linkage and network between carrier proteins and post de novo synthesis actors like desaturases (Zhao et al. 2019). Glyma.09G014000 is also linked to another desaturase gene, FAD8-1 (Glyma.01G120400) (Wang et al. 2016). Evidence also supports an interaction between different desaturases (**Figure 6. C**). FAD2-2 (Glyma.03g144500) interacts with ACPD (Glyma.02g138100) (**Figure 7**). ACPD is a stearoyl-[acyl-carrier-protein] 9-desaturase which converts stearoyl-ACP to oleoyl-ACP (Thambugala et al. 2013).

We detected interactions between Acetyl CoA carboxylase (ACCase) components and genes involved in malonyl-CoA biosynthesis. ACCase-B (Glyma.04G104900) is an acetyl-CoA carboxylase 1 gene with an interaction link to BCCP1 (Glyma.13g057400), a biotin carboxyl carrier protein 2 (**Figure 7**). BCCP1 also interacted with Glyma.09g277400, a 3-ketoacyl-acyl carrier protein synthase III (KAS III) (**Figure 7**). These interactions (between ACCase, BCCP1, and KAS III) are merely suggestive as we could not find experimental evidence in support in spite of having a STRING combined score > 0.9 in the PPI network. A high combined score suggests a high likelihood of interaction.

## Conclusions

We combined GWAS with a selective scan across the genome (GWASi) to capture the contribution of interactions between genomic regions that individually may not appear significantly associated with a trait. GWAS identified 251 independent mutations associated with the variation in fatty acid composition. We evaluated 55 genes encoding different acyl carriers, fatty acyl reductases, oxidases, dehydratases, elongases and desaturases which were located within 1000 kbp of significant associations. GWASi and functional enrichment analyses identified 7000+ significant interactions among variants and genes in their neighborhood pointing to the often neglected but key contribution of gene interactions in modifying phenotypes.

For fatty acid variability in peanut oil, we demonstrate that additive, epistatic, and pleiotropic causal variants underlie the proposed candidate genes controlling this complex trait. Our results speak to the complex nature of the genetic control of fatty acid composition in seed oils, beyond the well characterized mutations in ahFAD2 genes.

## Data Availability

Supplementary data is available at the National Ag Library for the genotype data: https://data.nal.usda.gov/dataset/data-genotypic-characterization-us-peanut-core-collection. All other supplementary data is included in the manuscript submission.

## Acknowledgments

Mention of trade names or commercial products in this publication is solely for the purpose of providing specific information and does not imply recommendation or endorsement by the U.S. Department of Agriculture. USDA is an equal opportunity provider and Employer. We thank David Fernández-Baca, Sudhansu Dash, Lori M. Lincoln, Ye Chu, Will Dezern, Roshan Kulkarni, and David J. Bertioli for their contributions to associated projects to genotype and phenotype the materials used in this study.

## Supplementary Material

**Figure S1. A.** Correlations among fatty acid components, **B.** LD decay across the genome and subgenomes.

**Figure S2.** Gene ontology (GO) enrichment terms of GWAS interacting SNPs. **A.** Molecular Function, **B.** Biological Process.

**Table S1.** Effect of population structure and population size on the quality of GWAS.

**Table S2.** GWAS significant regions and predicted candidate genes.

**Table S3.** Epistatic interactions contributing to variation in fatty acid concentration (CASSI, selective approach, P < 4.65E-08).

**Table S4.** Gene interactions identified via GWASi.

**Table S5.** Enrichment Analysis of GWAS interacting SNPs.

